# Islet macrophages drive islet vascular remodeling and compensatory hyperinsulinemia in the early stages of diabetes

**DOI:** 10.1101/584953

**Authors:** Manesh Chittezhath, Divya Gunaseelan, Xiaofeng Zheng, Riasat Hasan, Vanessa SY Tay, Seok Ting Lim, Xiaomeng Wang, Stefan Bornstein, Per-Olof Berggren, Bernhard Boehm, Christiane Ruedl, Yusuf Ali

## Abstract

β-cells respond to peripheral insulin resistance by increasing circulating insulin in early type-2 diabetes (T2D). Islet remodeling supports this compensation but the drivers of this process remain poorly understood. Infiltrating macrophages have been implicated in late stage T2D but relatively little is known on islet resident macrophages, especially in early T2D. We hypothesize that islet resident macrophages contribute to islet vascular remodeling and hyperinsulinemia, the failure of which results in a rapid progression to T2D. Using genetic and diet-induced models of compensatory hyperinsulinemia we show that its depletion significantly compromises islet remodeling in terms of size, vascular density and insulin secretion capacity. Depletion of islet macrophages reduces VEGF-A secretion from both human and mouse islets ex vivo and the impact of reduced VEGF-A functionally translates to delayed re-vascularization upon transplantation in vivo. Hence, we show a new role of islet resident macrophages in the context of early T2D and suggest that there is considerable utility in harnessing islet macrophages to promote islet remodeling and islet insulin secretion capacity.

**Highlights:**

- The compensatory hyperinsulinemic phase of type-2 diabetes is accompanied with significant pancreatic islet remodeling.
- *Bona fide* islet resident macrophages are increased during the diabetic compensation phase by largely *in situ* proliferation.
- Ablating macrophages severely compromises the islet remodeling process and exacerbates glycemic control *in vivo*.
- Mouse and human islet macrophages contribute VEGF-A to the islet environment.
- Specific removal of islet macrophages delays islet vascularization in compensatory hyperinsulinemic mice.

## Introduction

Even though insulin resistance is considered the major underlying feature of type 2 diabetes mellitus (T2D) pancreatic β-cell activity, and insulin insufficiency, remains central to the pathogenesis of this disease (1). β-cells initially respond to insulin resistance by inducing a compensatory homeostatic mechanism that results in hyperinsulinemia to prevent circulating glucose levels from going too high. This adaptive compensatory mechanism eventually fails due to pancreatic β-cell inactivity (either through dysfunction, dedifferentiation or apoptotic mechanisms) leading to the development of overt T2D (2). Although β-cell produce insulin, it is widely accepted that other neighboring islet cell types play an important role in supporting and mediating an appropriate β-cell response too. Prolonging the islet compensation process, by first gaining a better understanding of the paracrine factors that contribute to it, is one viable strategy to prevent insulin insufficiency and T2D pathogenesis.

Macrophages are the major immune cell type in pancreatic islets at steady state but their presence per islet remain low (3). Islet macrophages have long been reported to increase during prolonged obesity and in T2D but the functional consequences of increased islet macrophage have always been negative. These macrophages are often regarded as the major driver of islet inflammation and β-cell dysfunction and they thereby directly contribute to T2D development (4-7). On the other hand, islet macrophages have also been reported to be essential for β-cell formation during embryonic development and in experimental models of pancreatic β-cell regeneration, macrophages have been shown to support β-cell proliferation (8). Since macrophages are highly plastic with rapid phenotypic adaptations to the local environment, it is likely that islet macrophages play different roles at either end of the T2D pathogenesis spectrum (9-11).

Thus far, studies on islet macrophages have focused on the later stages of diabetic pathogenesis, when insulin secretion is compromised. For example, increased macrophage accumulation has been confirmed by pathological studies using pancreatic islets from T2D patients (12, 13) and from rodent models of obesity and type-2 diabetes (14, 15). In support of this, factors such as glucolipotoxicity, endotoxicity and islet amyloid deposition have been implicated in the islet pro-inflammatory macrophage recruitment stage that drive T2D pathogenesis (16-18).

Relatively little is known of islet macrophages in the early T2D phase of modest hyperglycemia and hyperinsulinemia. The latter is facilitated by islet remodeling and hypertrophy as well as β-cell hyperplasia. With evidence of macrophage involvement during embryonic pancreatic development and their importance in supporting β-cell replication in experimental rodent models of pancreatic regeneration, it is conceivable that islet macrophages may have a positive impact on islet and β-cell function (8, 19, 20). In addition, VEGFA-mediated β-cell expansion was shown to require increased macrophage presence to the islet micro-environment (21). These studies together hint at a possible beneficial role that islet macrophages may have. Although certain in the context of pancreatic development and β-cell regeneration following injury, macrophage involvement during the early stages of T2D and β-cell compensation remains nebulous (22).

We hypothesize that during islet compensation in early T2D, islet macrophages are required for islet vascular remodeling, increased insulin secretion and maintenance of postprandial glucose. To test this hypothesis we utilized two different islet compensation (genetic and diet-induced) mouse models and show that in both models there is a significant increase in islet macrophage numbers with evidence of intra-islet macrophage replication and expansion. Ablating these macrophages during the islet compensation phase with three different methods compromises the islet vascular remodeling process as evidenced primarily by reduced vascularization, maintained islet size (instead of increased as usually observed during islet compensation) and impaired insulin secretion. An exacerbation of glucose homeostasis was observed in mice devoid of islet macrophages and this is driven largely by reduced glucose-induced insulin secretion rather than a decrease in peripheral insulin sensitivity. Human islet macrophages, similar to mice, contribute VEGF-A and TNFα to the local milieu. Finally, we demonstrate that islet macrophages directly contribute to its remodeling by supporting islet vascularization, through secretion of VEGFA in the early stage of islet remodeling, suggesting its key role in supporting islet compensation in early T2D.

## Results

### Higher numbers of macrophages within islets of db/db mice

Macrophages are known to exist in pancreatic islets at steady state (23), however their role in early type-2 diabetes is still relatively unknown. Here, we looked specifically at islet compensation in young db/db mice (aged 16 weeks and less) with the hypothesis that islet macrophages contribute to the islet hypertrophy and β-cell hyperplasia observed in the mice at this age (collectively referred to as islet remodeling). The leptin-receptor deficient B6.BKS(D)-Lepr^db^/J mouse (db/db) has a modest but significant increase in fasting glucose levels, a significantly higher glucose excursion following intraperitoneal glucose injection, and a higher circulating insulin during both fasting- and glucose-stimulated states, compared with non-diabetic lean littermates (lean) (Supplemental Figure 1A-C). Islet size in db/db mice was consistently approximately 3.5 fold larger compared to lean mice (Supplemental Figure 1D-E), corroborating an earlier observation (24). These results are similar to the metabolic adaptations seen in pre-diabetic humans (1, 25, 26) making the db/db mouse an ideal model for studying mechanisms of islet compensation in early T2D.

We determined islet macrophage numbers during this T2D compensation process. Pancreatic sections from db/db and age-matched lean mice were immunolabelled with either F4/80 or CD68 to stain for macrophages (Figure 1B, C left panel). The percentage of F4/80^+^ positive cells were higher in the db/db islets compared to lean mice at 8, 12 and 16 weeks of age (Figure 1A, right panel). The observed increase in macrophage numbers in db/db mice was further corroborated with CD68^+^ immunofluorescence (Figure 1B right panel), suggesting that macrophages are present at higher numbers in db/db islets which is similar to a recent study showing increased islet macrophage numbers in genetically obese (ob/ob) mice(3). We further performed a sub-population analysis of freshly isolated and dissociated islet cells using flow cytometry with a panel of earlier-identified macrophage markers (23). In both experimental groups (lean versus db/db), isolated macrophages were mostly CD11b^+^, F4/80^hi^ and MHCII^hi^ and lacking the Ly6C expression (Figure 1C left panel). At 16 weeks of age, the number of macrophages relative to CD45^+^ cells was higher in islets obtained from db/db mice compared to islets collected from correspondent aged-matched lean controls (Figure 1C, right panel) suggesting that islet macrophages at 16 weeks of age were perhaps either resident self-renewing macrophages (23) or further infiltrating monocytes that subsequently lost their surface Ly6C expression upon differentiation. To further evaluate this, islets were immunolabelled with insulin, F4/80 and the proliferation marker Ki67. We do observe instances of Ki67^+^, F4/80^+^ cells suggesting islet macrophage cell replication, but the incidence is too low to account for the marked differences between 16 week db/db and lean islets (Figure 1D). These results suggest that, despite its low proportion among other islet cells, CD11b^+^F4/80^high^ macrophages are present at higher numbers in the compensating islet of db/db mice as a result of both *in situ* islet expansion and possibly monocyte infiltration.

**Figure 1.**
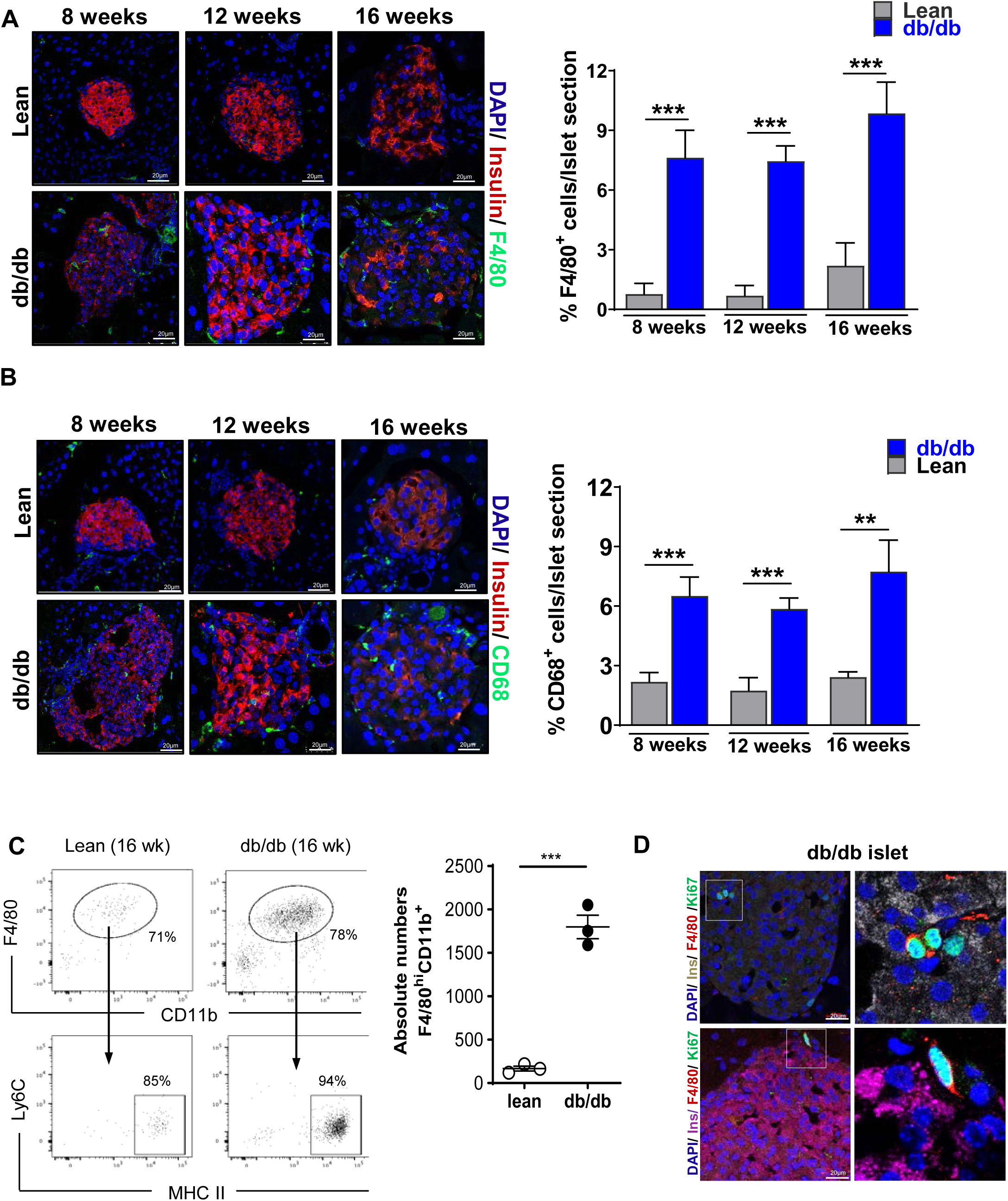
Increased macrophage numbers in the db/db compensating islet. Pancreatic islets from 8, 12 and 16 weeks old db/db and age-matched lean controls were immunolabelled with macrophage markers **(A).** F4/80, **(B).** CD68; Insulin (red), F4/80/ CD68 (green), DAPI (blue) left panel. Quantitation of CD68^+^ and F4/80^+^ cells (right panel). **C.** FACS analysis; Single cell dissociated Islets 16 weeks old db/db or age matched lean controls were analysed by flow cytometry by gating on single CD45+ cells. Islet-resident myeloid cells were cell surface positive for CD11b, and F4/80 and MHCII (Left panel). Quantification of absolute numbers of F4/80^hi^CD11b^+^ cells (right panel). Data is represented as mean ± SEM (n=4). Scale bar = 20 μm.P-values were calculated using unpaired Student’s t-test ***p<0.001; **p<0.005 Vs lean**. D.** Presence of proliferating macrophages in the pancreatic Islets. Two different islets (Top and Bottom Panels) are shown. Boxed area is enlarged in right panel. Insulin (Magenta/grey), F4/80 (red), proliferation marker Ki67 (green).

### Macrophages contribute to islet remodeling and whole body glucose homeostasis during compensation

To determine whether the observed increase in intra-islet macrophages is related to islet remodeling in db/db mice, we depleted macrophages using clodronate. Briefly, clodronate containing liposomes (50 mg/kg body weight) were administered i.p into 10 weeks old db/db (db/db clodronate) and age-matched control mice (lean clodronate) every 3 days for 2 weeks. Control group received injections of PBS-liposomes (db/db-PBS, Lean-PBS) at similar time points (Suppl. Figure 2A). Clodronate treatment resulted in a significant reduction of CD68^+^ positive macrophages in db/db islets compared to db/db islets in the PBS-treated group (9.4% in db/db-PBS Vs 1.4% in db/db-clo) (Figure 2A left and middle panel). Lean clodronate-treated mice also showed a modest but insignificant reduction in islet CD68^+^ cell numbers compared to lean-PBS and the lack of significance perhaps reflects the relatively low number of macrophages in lean islets to begin with (Figure 2A middle panel). Noteworthy, the extent of reduction in islet macrophage was not seen in other tissues such as adipose tissue of clodronate-treated db/db compared to PBS treated db/db because islets have proportionately few macrophages compared to fat (Supplemental Figure 2B). Although clodronate treatment resulted in an elevated insulin intolerance, this difference was not statistically significant when compared to PBS liposome-treated db/db mice (Supplemental Figure 2C). There is a trend of reduced insulin sensitivity when db/db mice were treated with clodronate but this is against what is reported in literature, where clodronate treatment was shown to improve glycemic parameters in treated mice (14).

**Figure 2.**
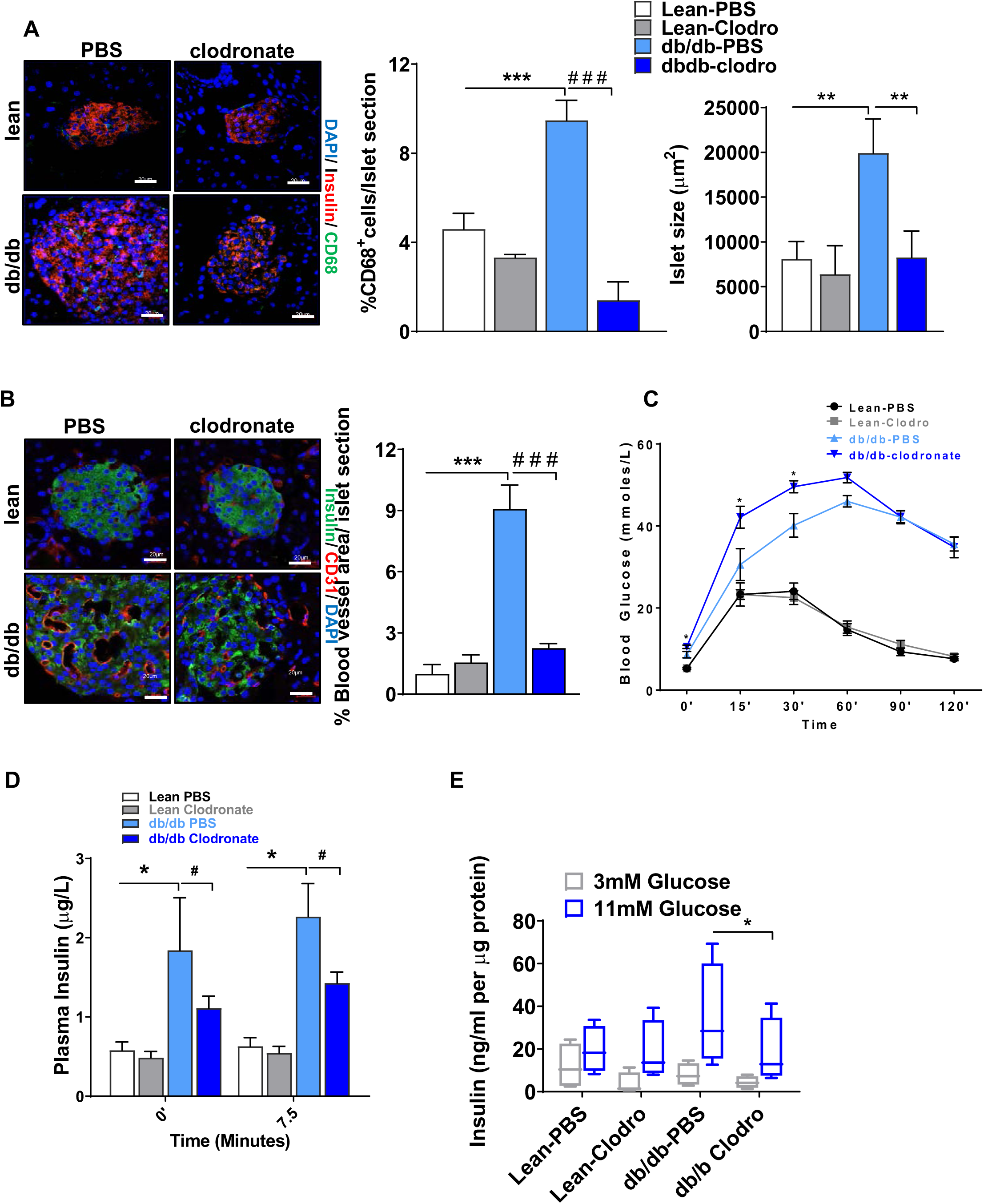
Islet macrophages contribute to islet remodeling and glucose homeostasis during compensation. **A.** Representative immunofluorescence image showing depletion of CD68^+^ macrophages after clodronate treatment in islet (left panel) and quantification of CD68^+^ macrophages (middle panel) and islet size (right panel). Insulin (red), CD68 (green), DAPI (blue). **B.** Representative image (left panel) showing effect of clodronate treatment on islet vasculature. Insulin (green), endothelial marker CD31 (red) and DAPI (blue). Quantitation of blood vessel area (right panel). **C.** Intraperitoneal glucose tolerance test (IPGTT) in lean and db/db mice treated with either PBS-liposomes or Clodronate-liposomes. **D.** Level of plasma insulin before and after intraperitoneal loading with 2g D-glucose/kg on 12-week-old db/db or lean controls or treated with clodronate after overnight fasting. **E.** Insulin secretion from isolated clodronate treated pancreatic islets *ex vivo*. Insulin secretion in response to the indicated secretagogues. Values are expressed in nanograms of insulin/ml per µg protein. All data in this figure are the means ± SEM (scale bar, 20 μm). P-values were calculated using One Way ANOVA with Tukey’s multiple comparisons test (**a**, **b**) and unpaired Student’s t-test (**c**, **d**, **e**) ***p<0.001, **p<0.005 *p<0.05 Vs lean-PBS; ^###^p<0.001; ^#^p<0.05 Vs db/db clodronate.

We next determined the effect of islet macrophage depletion on islet morphology, vasculature and function. Clodronate treatment resulted in reduced db/db islet size with a concomitant decrease in beta cell^+^ area, especially in clodronate treated db/db mice (Figure 2A right panel). Islet vasculature area (CD31^+^area) as a percentage of total islet area was significantly reduced in clodronate-treated db/db islets compared to db/db-PBS control islets (Figure 2B). When given an i.p glucose load, db/db clodronate mice had a significantly higher glucose excursion particularly at the early time points of 15 and 30 min post glucose load (Figure 2C). Compensatory hyperinsulinemia is clearly seen in db/db mice. However, when treated with clodronate, both fasted and stimulated insulin were significantly reduced in the db/db clodronate group (Figure 2D). Consistent with this, islet macrophage depletion resulted in a modest but significant reduction in insulin secretion when islets were stimulated with high glucose *ex vivo* (Figure 2E). Taken together, these data suggests that macrophages contribute to islet remodeling and compensatory hypersulinemia in the db/db mice. By reducing macrophages, including islet macrophages, islet remodeling is curtailed and compensatory hyperinsulinemia is markedly compromised making db/db mice more severely diabetic.

### Human and mouse islet macrophages contribute VEGF-A and inflammatory cytokines to the local milieu

To understand the molecular drivers underpinning islet remodeling, especially in the absence of islet macrophages, we measured cytokines in the supernatant of overnight islet cultures. Consistent with the effect of islet macrophage depletion on islet morphology and vasculature, our islet cytokine release analysis showed first and foremost a significant reduction in islet-derived VEGFA, a known regulator of islet vascular development and vascular homeostasis (21, 27-29) (Figure 3A). *Ex vivo* macrophage depletion also resulted in a significant reduction in pro-inflammatory cytokines release such as IL-6, IP-10 and G-CSF (Figure 3A). These cytokines are those that can be detected in the islet culture media and they have been shown to be produced by macrophages suggesting that in the compensation phase of T2D, macrophages release pro-inflammatory cytokines at a level that is sufficient to stimulate the islet remodeling process (Figure 3A). The pro-inflammatory contribution of macrophages to the islet milieu has been previously reported (30). An *ex vivo* depletion of macrophages from both lean and 12-week db/db mouse islets did not significantly alter β-cell signature genes after 24-hours in culture suggesting that β-cells themselves were perhaps not directly impacted by the loss of macrophages (Figure 3B). Next, we analyzed the presence of macrophages in human islets using the best available human microglia/macrophage-specific marker Iba-1. Freshly isolated human islets contain a similar number of macrophages as mouse islets and the Iba-1 positivity in islets was significantly reduced with *ex vivo* clodronate treatment (Figure 3C, D). We repeated these experiments using three different batches of human islets that were obtained from non-diabetic donors. Human islets at steady state secrete VEGF-A and a number of measurable pro-inflammatory cytokines namely TNFα, IL-6 and IL-10 *ex vivo* corroborating an earlier observation (Figure 3E) (31). Next, *ex vivo* depletion of macrophages from human islets resulted in a significant reduction of both VEGF-A and TNFα, suggesting that these two paracrine factors are largely derived from macrophages within the islet. IL-10 was modestly reduced but its relatively low abundance may account for the lack of significance. Unlike mouse islets however, IL-6 was increased by almost 3-fold in non-diabetic human islets devoid of macrophages suggesting that IL-6 in human islets do not originate from macrophages and that islet macrophage removal induces the further release of IL-6, which have been reported to be secreted by β-cells (32). However, the lack of human islet macrophages, and increase in IL-6 which has been shown to be protective to β-cells, do not seem to significantly affect β-cells as there are no significant changes in human β-cell signature genes following clodronate treatment (Figure 3F). This lack of significance also suggests that islet macrophages indirectly impact human β-cells through another cell type that is heavily involved in islet remodeling.

**Figure 3.**
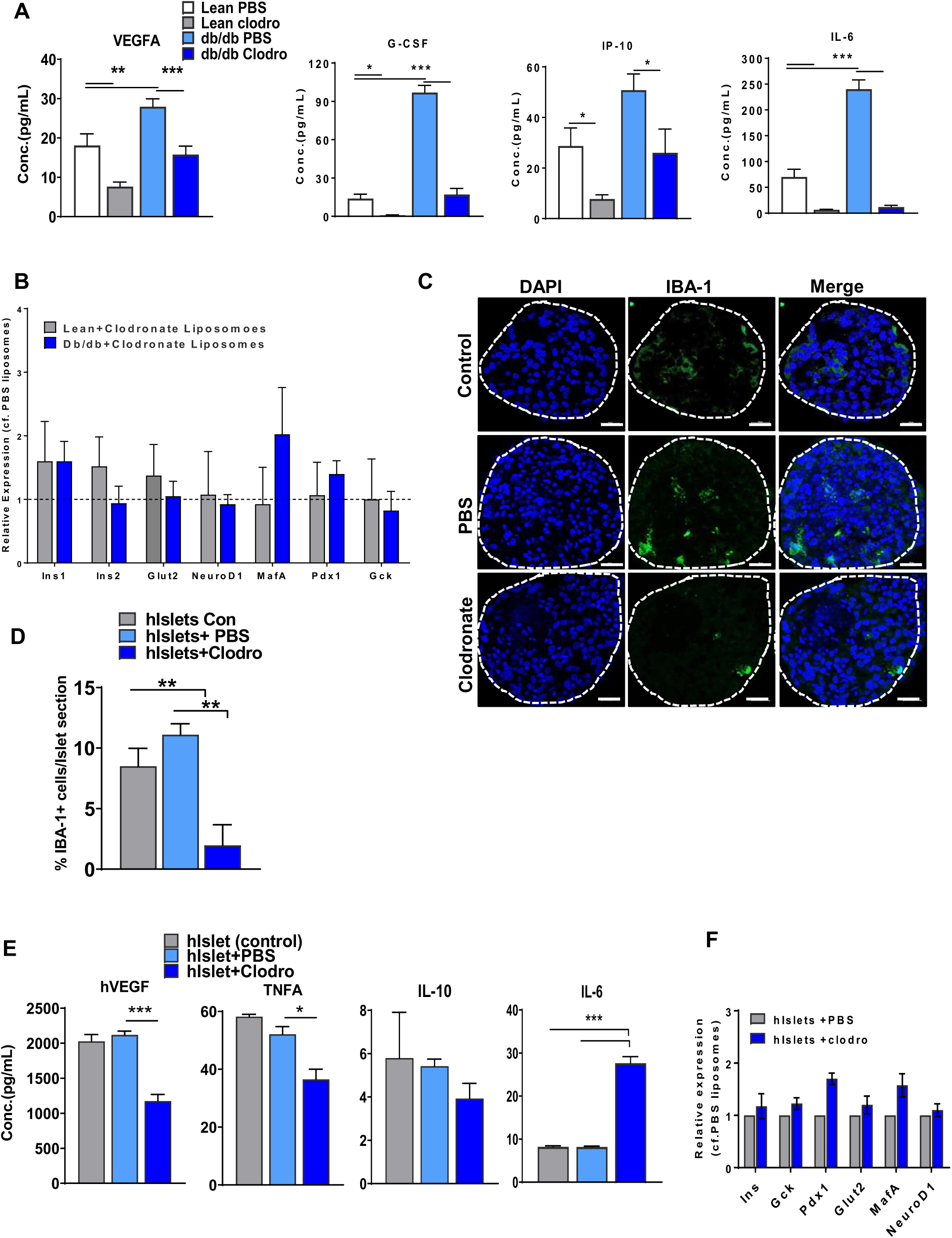
Mouse and human islets secrete VEGF-A and other cytokines. **A.** Macrophage depletion by clodronate treatment alters the cytokine profile of pancreatic islets from db/db and lean mice *ex vivo*: Cytokine secretion was measured on a Bio-Plex 200 reader and data analyzed by Bio-Plex Data Pro(tm) Software as mean fluorescent intensity (MFI). All samples were run in duplicates, N=5. **B.** qPCR analysis of β-cell specific genes in lean and db/db islets after clodronate treatment. **C.** Presence of macrophages in human islets. Representative immunofluorescence image with macrophage marker IBA-1 (green), DAPI (Blue). Dotted line indicate the islet area. **D.** Quantification of IBA^+^ cells in the islet after clodronate treatment. **E.** Cytokine profile of human pancreatic islets after *ex vivo* clodronate treatment using Milliplex MAP kit. **F.** qPCR analysis of β-cell specific genes in human islets after clodronate treatment. All data in this figure are the means ± SEM (scale bar, 20 μm). P-values were calculated using One Way ANOVA withTukey’s multiple comparisons test ***p<0.001, **p<0.005 *p<0.05.

### Macrophage ablation reduces islet remodeling and insulin secretion in High Fat Diet-fed CD169-DTR mice

Thus far, we have relied on one mouse model to suggest that islet macrophages are implicated in the islet remodeling process of compensatory hyperinsulinemia. To confirm our findings, we determined macrophage involvement in a diet-induced model of compensatory hyperinsulinemia and subjected this model to two different macrophage ablation protocols. Male C57BL/6 mice were fed high fat diet (HFD) or calorifically-matched low fat diet (LFD) for just an acute period where pancreatic tissues were harvested after 3, 7, 14 and 21 days of feeding. Pancreatic sections were stained for F4/80 and CD68 to quantify islet macrophages across this acute time period. We observed an increase in F4/80^+^ cells at 21 days with the percentage of islet macrophages increasing significantly (by approximately 3-fold) compared to calorie-matched LFD-fed mice (Figure 4A, 4B). Similarly, we observed a significant increase in the CD68^+^ macrophage population in the islets at day 21 of HFD feeding (Supplemental Figure 3A) but not during earlier time points suggesting that 21 days of HFD is minimally required for significant changes in islet macrophage numbers to occur. This is contrast to an earlier study that reported increase in islet macrophages as early as 7 days after HFD feeding, but their controls were not calorifically matched and thus may explain the difference in results obtained(33).

**Figure 4.**
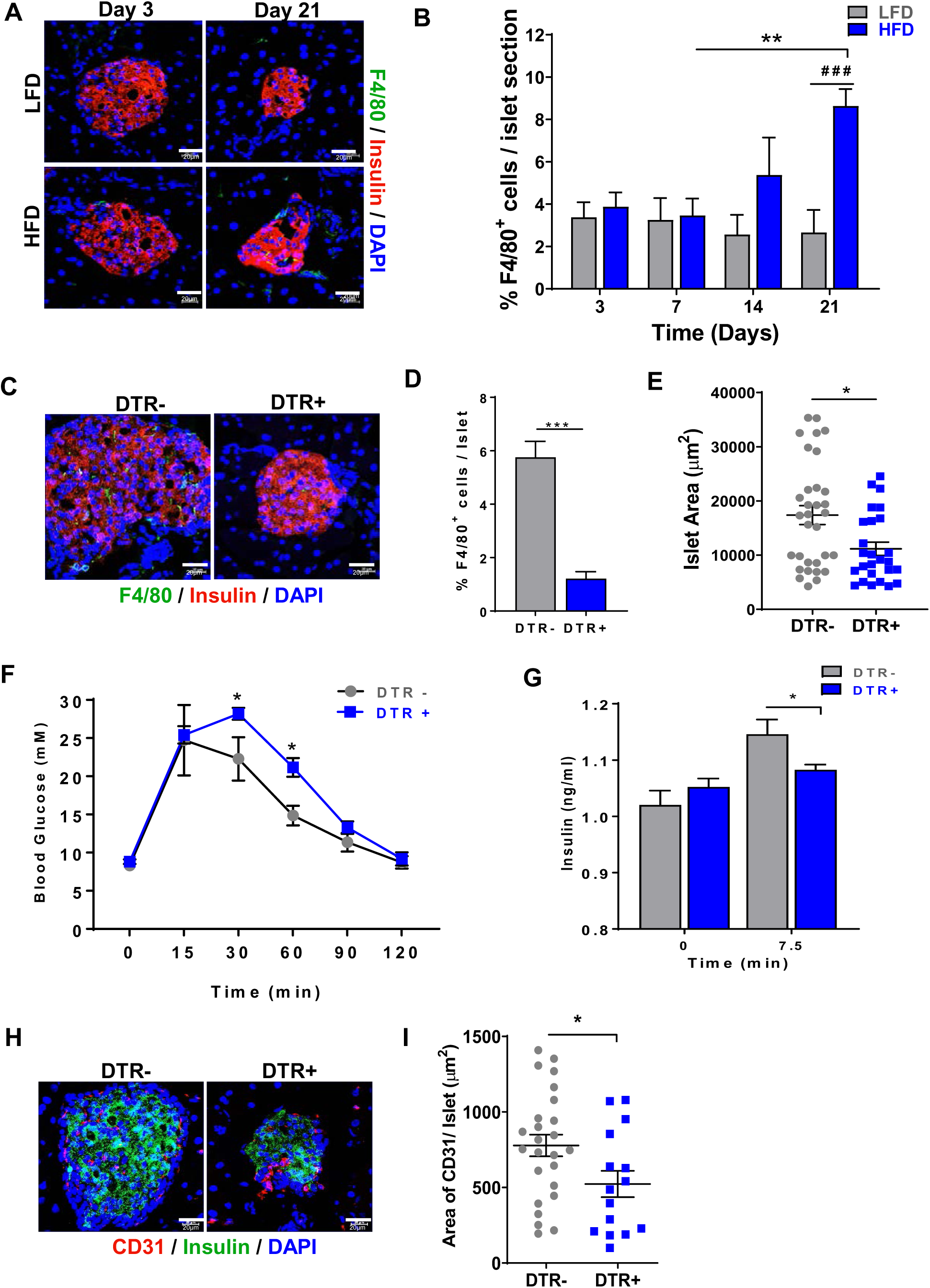
Effect of macrophage depletion on islet remodeling and function in CD169-DTR mouse model. **A.** Representative image showing F4/80^+^ macrophages in HFD fed mice compared to LFD fed mice and, **B.** quantification of F4/80^+^ cells/islet (right panel). Insulin (red), F4/80 (green), DAPI (blue). **C.** Representative image showing the depletion of F4/80^+^ macrophages in islets of CD169-DTR^+^ mice after DT injection. Insulin (red), F4/80 (green), DAPI (blue) (left panel) and **D**. quantification of F4/80^+^ cells/islet. **E.** Islet area of CD169-DTR^−^ and CD169-DTR^+^ mice after DT injection. **F.** Intraperitoneal glucose tolerance test (IPGTT) after overnight fasting. **G.** Plasma Insulin before and 7.5 min after glucose challenge. **H.** Representative image showing effect of macrophage depletion on islet vasculature in CD169-DTR^+^ mice. Insulin (green), endothelial marker CD31 (red) and DAPI (blue). **I.** Quantitation of blood vessel area based on CD31^+^ area per islet. All data in this figure are the means ± SEM (scale bar, 20 μm). P-values were calculated using unpaired Student’s t-test ***p<0.001, **p<0.005, *p<0.05, ^###^p<0.001 Vs lean.

In our hands, a significant increase in body weight was noted in HFD-fed mice compared to the calorifically matched LFD-fed mice at day 20 (Supplemental Figure 3B). In addition, the increase in macrophage number in HFD-fed islets coincided with an increase in islet size at day 21 (Supplemental Figure 3C). There was a modest but significantly higher fasting blood glucose in HFD on day 21 compared to LFD-fed mice and a glucose challenge significantly increased the glucose excursion following a glucose load in the HFD-fed mice (Supplemental Figure 3D left and middle panel). Fasting insulin was significantly increased after HFD feeding from day 10 to 20 (Supplemental Figure 3D right panel) suggesting that a 21-day HFD diet induces slight but significant hyperinsulinemia (compared to day 10 HFD-fed mice). Correspondingly, we observed an increase in islet vasculature at day 21 in the HFD-fed mice (Supplemental Figure 3E). Taken together these results suggest that a 3-week HFD treatment induced compensatory islet hyperinsulinemia and islet remodeling with observed increases in islet size, vasculature and islet macrophage numbers.

We next proceeded to determine the effect of islet macrophage depletion on islet remodeling and compensatory hyperinsulinemia in 21 day HFD-fed mice. We used CD169 diphtheria toxin receptor (DTR) mouse model where injection of diphtheria toxin (DT) administration specifically ablates the CD169 (sialoadhesin) expressing F4/80^hi^ resident macrophages(34). Both DTR^+^ and DTR^−^ control mice were fed with HFD with DT administration every 3 days for 21 days (Supplemental Figure 4A). Measurements of body weight (Supplemental Figure 4B), IPGTT and serum insulin (Supplemental Figure 4C) were made to determine the effect of macrophage depletion on glucose homeostasis in HFD-fed mice. DT Injection resulted in a significant depletion of F4/80^+^ positive macrophages in DTR^+^ mice on HFD compared to DTR^−^ mice at day 21 (Figure 4C, 4D). Consistent with our db/db mouse-chlodronate studies, DT treatment resulted in a significant reduction in islet area (Figure 4E). Macrophage depletion in HFD-treated DTR^+^ mice resulted in a significant decrease in body weight compared to the DTR^−^ mice on HFD at day 21 (Supplemental Figure 4B) even though there were no significant changes in fasting blood glucose levels between HFD-fed DTR^+^ mice and HFD-fed DTR^−^ mice (Supplemental Figure 4C). The reduction in body weight of HFD mice has been reported in HFD mice treated with clodronate for 4 weeks, suggesting that systemic ablation of macrophages do have an effect on HFD-mediated weight gain(35). There was a modest glucose intolerance in HFD treated DTR^+^ mice which was significant at 30 and 60min after a glucose challenge compared to HFD-fed DTR^−^ mice (Figure 4F). Although there was no difference in fasting serum insulin levels in both DTR^+^ and DTR^−^ mice, the endogenous insulin secretion upon glucose stimulation was significantly lower in the HFD fed DTR^+^ mice compared to HFD-fed DTR^−^ mice at day 21 (Figure 4G). There was a decrease in insulin stimulated glucose clearance in DTR^+^ mice on HFD at day 21 which was significant at 60 and 90 min after an insulin challenge (ITT) suggesting a decrease in peripheral insulin sensitivity in DT-treated HFD mice (Supplemental Figure 4D). Depletion of macrophages in DTR^+^ mice resulted in a reduction in the blood vessel area as shown by CD31 staining, compared to DTR^−^ mice (Figure 4H-I). Overall, depletion of macrophages and HFD feeding for an acute term of 21 days affected the body weight of the mice but changes were also noted in islet size, β-cell number and islet vasculature which may explain the reduced glucose-stimulated insulin secretion and the impaired glucose tolerance in these mice.

### Macrophage depletion reduces islet remodeling and insulin secretion in CSF-1R treated HFD-fed mice

To confirm macrophage involvement in islet remodeling and function during HFD-fed compensation in transgenic mice (CD169-DTR), we repeated experiments in BL6/N mice treated with a monoclonal antibody against CSF-1R (AFS98). CSF-1R anti-body was previously shown to significantly reduce islet macrophage numbers in type-1 diabetic islets (36). We tested whether inhibition of CSF-1R (CSF-1Rα) could prevent macrophage dependent islet remodeling and vascular development in HFD-fed mice. Intraperitoneal administration of AFS98 (CSF1-R α, 2mg/kg body weight; i.p; Day1, Day10 and day 20) (Figure 5A) led to ∽80% reduction of the islet macrophages in HFD-fed mice at day 21 (Figure 5B-C). There was a significant reduction in the islet size in CSF-1R antibody treated mice similar to CD169-DTR mice (Figue 5D), with no change in body weight (data not shown). Glucose excursion was trending higher in the AFS98-treated mice with a significant difference noted at the 120 min time-point (Figure 5E). However, insulin sensitivity were unaltered between antibody treated and control groups (Figure 5F). The modest but significant increase in glucose excursion was perhaps due to plasma insulin levels being significantly reduced 7.5 min after glucose challenge in the AFS98 treated mice (Figure 5G). Consistent with the earlier macrophage depletion experiments, there was a marked decrease in islet vascular density after AFS98 treatment (Figure 5H-I) suggesting that macrophage inhibition through CSF-1R antagonism severely impairs endothelial cell growth within islets which in turn impacts on islet remodeling, insulin release and glucose homeostasis.

**Figure 5.**
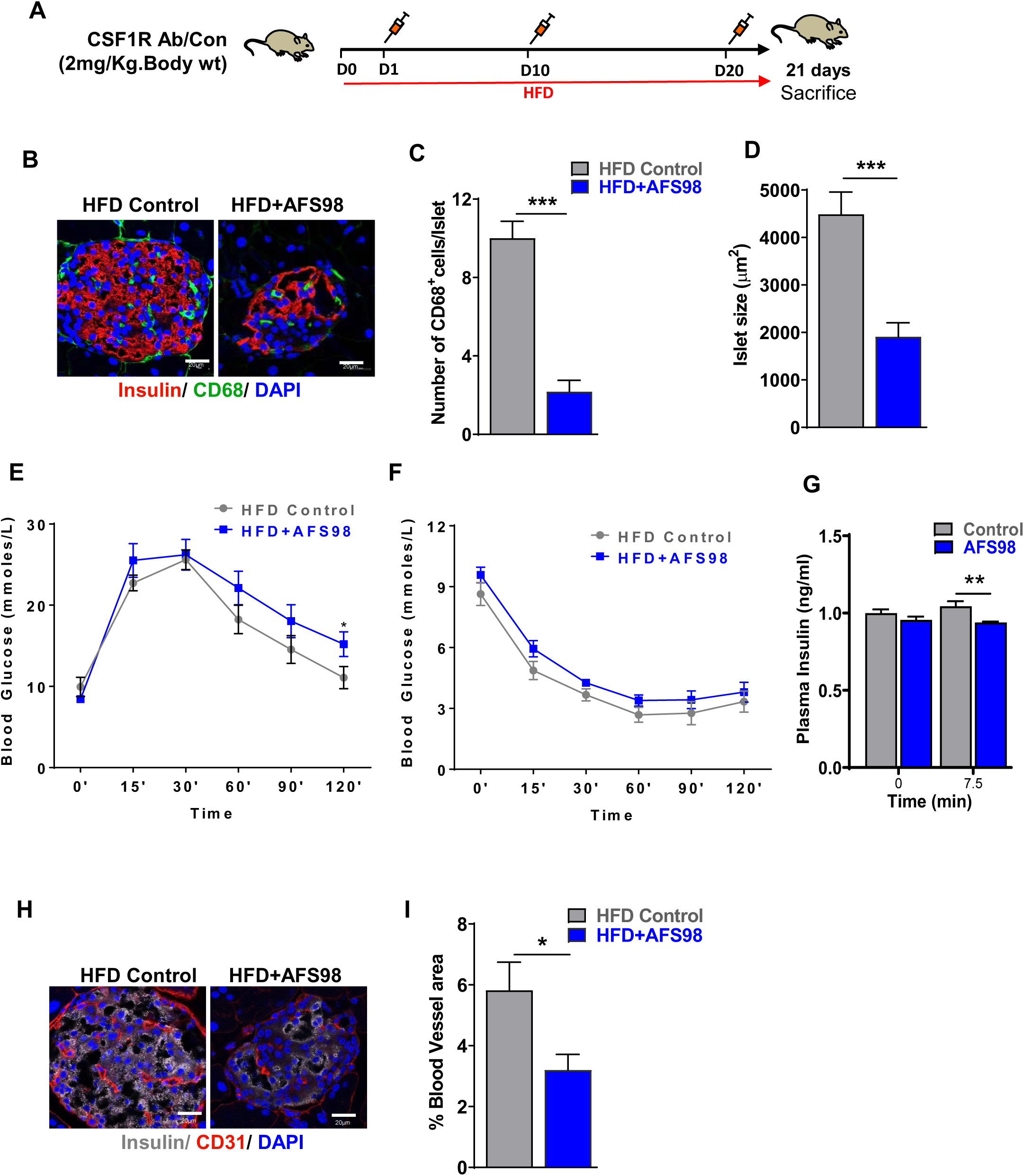
Macrophage depletion reduces islet vascular remodeling and insulin secretion in CSF-1R antibody treated HFD-fed mice**. A.** Schematic representation of the experimental design for macrophage depletion in CSF-1R antibody treated HFD-fed mice. **B.** Representative image showing macrophages depletion in CSF-1R antibody treated HFD fed mice compared to control Insulin (red), CD68 (green), DAPI (blue) and **C**. quantification of CD68^+^cells/islet. **D.** measurement of islet size **E.** Intraperitoneal glucose tolerance test (IPGTT), **F.** Insulin tolerance test (ITT), **G.** Plasma insulin before and after glucose injection in HFD-control and HFD-AFS98 treated mice. **H.** Representative image showing effect of CSF-1R ab treatment on islet vasculature. Insulin (grey), endothelial marker CD31 (red) and DAPI (blue). **I.** Quantitation of blood vessel area (right panel). Scale bar is 20µm. P-values were calculated using students t-test. **p<0.001, *p<0.05.

### Ablation of islet macrophages delays islet revascularization after transplantation

Through whole body macrophage depletion experiments in two different mouse models of islet compensation we show that macrophages are required for vascular islet density increase and islet hyperplasia. We showed that when treated with clodronate ex vivo, mouse and human islets have significantly lower levels of secreted VEGF-A. To functionally determine the impact of macrophage-produced VEGF-A and to show that islet macrophages are directly responsible for islet re-vascularization and islet vascular density, 21 day HFD islets were isolated and treated ex vivo with clodronate (or PBS containing liposomes) overnight before transplantation back into HFD fed recipient mice. To visually follow islet engraftment, the transplantation was done in the anterior chamber of the eye (ACE). Islet re-vascularization was imaged by retro-orbital injection of fluorescein isothiocyanate (FITC)-dextran (37). In control animals studied, islet re-vascularization started between 3 and 10 days after islet transplantation(38). Islet re-vascularization was initiated by large-diameter vessels, followed by progressive vessel branching and reduced overall vessel size as reported before(39) (Figure 6A). There was a significant delay in islet revascularization and significant decrease in the vessel density in macrophage-depleted islets compared to macrophage intact islets as measured by tail vein injection of FITC-labelled dextran to label all blood capillaries (Figure 6B-C). After 4 weeks post-transplantation, the eyes were removed and stained for F4/80, CD31 and insulin. There was a notable absence of F4/80 signal in the engrafted clodronate-treated islet suggesting that islet macrophages do not recover following ex vivo depletion (Figure 6D). There was considerable less CD31^+^ staining in engrafted clodronate-treated islets compared to control corroborating observations obtained from the intravital islet blood capillary data (Figure 6D). Taken together, these data suggest that islet macrophages, through VEGF-A production to the local islet milieu, have a direct impact on islet vascular growth with an impact on the ability of islets to undergo compensation to combat T2D. Without macrophages, islets do still get vascularized albeit at a slower rate because of reduced islet VEGF-A secretion levels compared to islets with macrophages. To further confirm the effect of macrophage depletion on blood vessel development, we investigated the effects of conditioned media from *ex vivo* macrophage depleted/ intact human islets HPaMEC angiogenesis. Using the *in-vitro* Matrigel tube formation assay, HPaMEC exposed to macrophage-free (clodronate liposome) human islets conditioned media had a marginal but consistent reduction in the number of meshes, segments, total tube length, and vessel branching intervals compared to macrophage-intact (PBS-liposome) islet conditioned media (Supplemental Figure 6A-B). This suggests that the impact of islet macrophage loss compromises the pro-angiogenic milieu.

**Figure 6.**
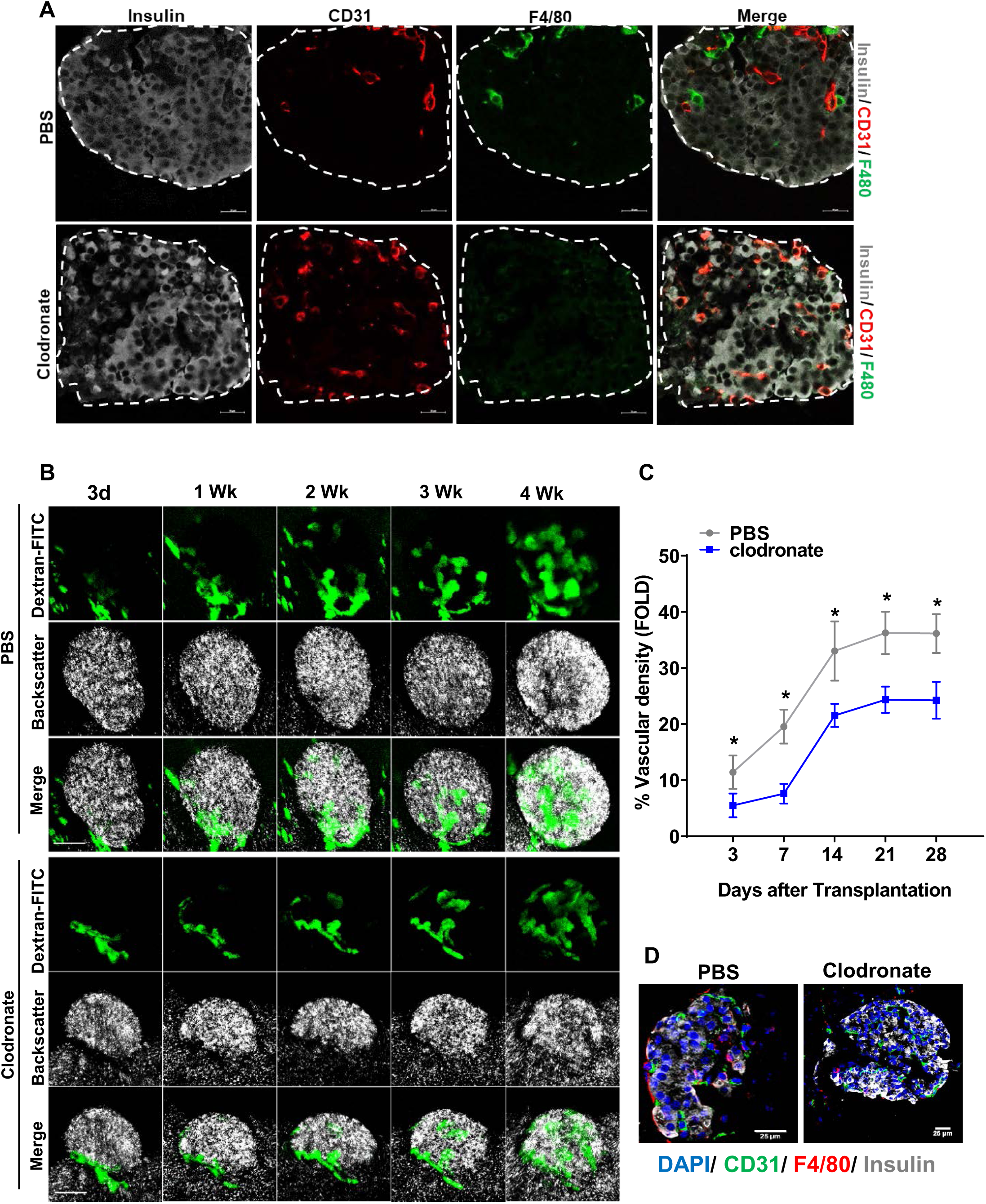
Selective ablation of islet macrophages affects islet revascularization upon transplantation into 21-day HFD-fed mice. **A.** Representative images of islets harvested from 21-day HFD fed mice treated with clodronate-liposomes or PBS-liposomes and stained for insulin (grey), endothelial marker CD31 (red) and F4/80 (green) doted lines demarcate islet. **B.** Longitudinal confocal imaging of the re-vascularization process of islets engrafted in the eye of HFD fed mice on indicated time intervals over a period of one month. Vessels (green) were stained in vivo with FITC-labelled 150-kDa dextran. Backscatter shown in white. **C.** Measurements of the vascular density of engrafted islets over time. Scale bar = 20µm. P-values were calculated using students t-test. *p<0.05. **D.** Representative image showing immuno-stained sections of islets 4 weeks post-transplantation. Frozen enucleated eyes containing the islets were sectioned and stained with endothelial marker CD31 (Green), F4/80 (Red), Insulin (Grey) and nuclei were counterstained with DAPI (Blue). Scale bar: 25 µm.

## Discussion

Islet macrophages have long been labelled as pathogenic contributing to islet inflammation and β-cell dysfunction. However none so far have addressed why, if so damaging, do these macrophages remain present in the islet in the first place? In the present study we decided to investigate the role of islet macrophages during the, often neglected, early T2D phase of islet compensation. In young genetically obese db/db mice (8-16 weeks) with a strong islet compensation phenotype, islet remodeling and hyperplasia (40) is correlated with higher numbers of macrophages within islets. The compensation process in db/db mice has been shown to start as early at 4 weeks, peak at 12 weeks and wane after 24 weeks of age (41). Our islet macrophage observations during the 8-16 week of age period are therefore representative of peaking islet compensation in these mice. A chemical-induced depletion of macrophages starting at 10 weeks of age for two weeks in db/db mice significantly reduced islet vascular density, islet hyperplasia, glucose-stimulated insulin secretion and glucose tolerance. These observations suggest that earlier gains of islet compensation were nullified when macrophages were removed. All these metabolic changes were also noted, albeit to a lesser extent in an acute HFD model of compensatory diabetes, where changes in islet remodeling were not as robust as those seen in young db/db mice. Similarly, upon macrophage ablation either genetically (CD169-DTR) or antibody-mediated (CSF1Rα treatment), led to compromised islet hyperplasia, reduced islet vascular density and blunted glucose-stimulated insulin response with a modest but significant impact on glucose tolerance in acute HFD mice. To date, there is no possible way to specifically ablate islet macrophages within the pancreas while leaving macrophages in other tissues intact. To overcome this limitation we utilized the anterior chamber of the eye (ACE) transplantation technique and studied how islets with and without macrophages behave upon transplantation into a HFD-fed mouse. Islets devoid of macrophages (*ex vivo* clodronate liposome treatment) were slower to revascularize compared to macrophage-intact transplanted islets. The delay in vascularization is a functional consequence of reduced islet VEGF-A levels, an angiogenesis factor that is produced, albeit not exclusively, by islet macrophages. Our results show for the first time that islet macrophage VEGF-A contribute to the increased VEGF-A islet milieu to induce islet vascular remodeling, β-cell hyperinsulinemia and the whole body maintenance of glucose sensitivity in the early phase of T2D.

We also show that the increased islet macrophage numbers were CD11b^+^, F4/80^hi^, MHCIII^+^, but lacking Ly6C expression. This islet macrophage phenotype is similarly described in both healthy and obese mice (3, 23). The relatively low expression of Ly6C on these cells indicate that these cells were either islet resident macrophages with a self-renewal capacity, or Ly6C^+^ inflammatory monocytes that subsequently shed the Ly6C positivity upon infiltrating the pancreatic islet. We directly observed Ki67^+^F4/80^+^ cells in db/db islets suggesting *in situ* islet resident macrophage replication, but it remains unclear whether replicating macrophages fully account for the observed increase in macrophage numbers in the compensating islet. Another possible mechanism for the increased Ly6C^−^ macrophages, as noted recently in an obesity model is the movement of peri-islet macrophages into the islet microenvironment. However, macrophage recruitment either from islet periphery or from the monocytic pool has been recently discounted through elegant observations using tagged monocytes in an obese T2D model (3). The expression of MHCII on islet macrophages suggest their role as antigen presenting cells which is perhaps critical for the clearance of debris and dead cells during tissue remodeling process to maintain tissue homeostasis.

Through clodronate-liposome based macrophage depletion experiments, we were able to determine the role of islet macrophages in islet compensatory hyperinsulinemia. Similar to earlier studies showing that overall islet morphology is perturbed and islet size is reduced by macrophage deficiency in csf-1deficient *(op/op)* mice during development (19), we found that compensatory islet remodeling and islet hyperplasia was significantly compromised in macrophage depleted db/db islets compared to islets from PBS treated db/db mice. Depletion of macrophages also led to a severe reduction of intra-islet vascular density, as evident by reduced numbers of CD31^+^ cells. This observation was consistent with a reduction in VEGF-A secretion, a principal regulator of islet vascular development (28) as well as an important macrophage chemoattractant in the islet culture supernatant from clodronate treated db/db mice. Studies have shown that, VEGF-A produced with in the pancreatic islets is essential for islet vascularization, revascularization, and function (28) suggesting that during islet compensation macrophage-derived VEGF-A, which is present in both mouse and human islets, contributes to islet vascularization. The functional consequence of reduced VEGF-A was further verified using the *in vivo* ACE-transplanted islet imaging technology. The ablation of mouse islet macrophages prior to transplantation significantly delayed islet revascularization and this led to a significant decrease in the vessel area in the macrophage-depleted islets compared to control transplanted islets. In line with these studies macrophage depletion compromised islet compensation and resulted in reduced insulin secretion from db/db islets treated with clodronate. This is particularly important, as the increased blood glucose level and insulin resistance in the diabetic mice demand increased insulin secretion from the pancreatic β-cells to maintain the blood glucose homeostasis and to maintain a pre-diabetic state. Key roles played by macrophages in islet remodeling during compensation has been further verified and confirmed in CD169-DTR as well as CSF1-R antibody-treatment in acute HFD mice.

We found the presence of macrophages in non-diabetic human islets and *ex vivo* culture analysis suggests that, similar to mouse islets, a number of proinflammatory cytokines are secreted from islets. We successfully depleted macrophages from human islets using clodronate *ex vivo* and observed that VEGF-A along with TNFα are significantly reduced. Unlike mouse islets, IL-6 was increased in macrophage depleted human islets and this observation is particularly intriguing because a number of studies allude to the β-cells protective effects of IL-6 (42-44). Nonetheless, similar to mouse islets, there was no significant change in the β-cell gene expression profile in human islets when macrophages were depleted further supporting the notion that islet macrophages have a greater impact on islet vasculature more so compared to a direct impact on β-cells. Further studies are warranted to identify whether the functional consequences of macrophage depletion in human islets results in reduced revascularization, as seen with mouse islets.

It is becoming increasingly clear that a pro-inflammatory environment in obese pancreatic islets contribute to β-cell dysfunction and insulin insufficiency leading to frank T2D (45, 46). The evidence for inflammation in islets (insulitis) during T2D is supported by increased infiltration of immune cells observed in islets of T2D patients and is corroborated by increased levels of secreted pro-inflammatory cytokines and chemokines from these islets (12, 47, 48). However, macrophages are major immune cell infiltrate shown to be critical for β-cell mass regeneration after injury by partial duct ligation (8) and shown to regulate VEGFA-mediated endothelial cell expansion and β-cell mass regeneration either directly or in cooperation with intra-islet endothelial cells (28). In a partial pancreatic ductal ligation (PDL) model, macrophage depletion by clodronate revealed the critical role of F4/80+ macrophages recruited to islets for β cell proliferation and these macrophages were found to be alternatively activated M2-like phenotype and contributed to the microenvironment necessary for β cell proliferation (8). Macrophages are also shown to be essential for CTGF-mediated adult β-cell proliferation in the setting of diphtheria toxin mediated partial β-cell ablation via increasing β-cell immaturity and shortening the replicative refractory period (49) A recent paper suggests that islet macrophages may act as islet sentinels and functionally react to ATP released by β-cells to promote a stable islet composition and glucose homeostasis at steady state (50). Taken together, a clearer picture on how the islet microenvironment shapes the phenotype of islet macrophages is emerging. However, our data coupled with recent published work also show islet macrophages themselves can directly impact islet function and whole body glucose homeostasis. Again, the nature of its role (positive for islet compensation or perhaps negative for chronic obesity) is context-dependent and involves the macrophage/ vasculature / β-cell axis.

Our data suggest a dual role played by islet macrophages at either end of the T2D pathogenesis spectrum. Macrophages are beneficial in early compensatory diabetes but this benefit is perhaps reversed with persistent and heightened pro-inflammation which then drives islet decompensation and β-cell loss. Our results point to a previous underappreciated beneficial role of islet macrophages in early compensatory T2D. Taken together, increased islet macrophage numbers in early T2D is required for increased islet blood vessel size and numbers and without islet macrophages the ability of islets to enhance insulin secretion is diminished. Our study highlights the significance of understanding the importance of islet macrophages in promoting islet vasculature growth, islet remodeling and hyperinsulinemia to compensate for insulin resistance. Apart from VEGF-A, these macrophages secrete a number of pro-inflammatory cytokines that is perhaps important for its own expansion within the islet. Our results suggest that the pre-diabetic state can be maintained with an appropriate level of pro-inflammatory islet signaling but yet again, should this inflammatory balance be upset islets will fail to compensate leading to a state of insulin insufficiency. Hence, a detailed functional characterization of islet macrophages during the different stages of diabetes progression may hold the key to determining the most appropriate therapeutic intervention to maintain islet integrity, β-cell function and glucose homeostasis during T2D pathogenesis.

## Acknowledgements

This work was supported by the Ministry of Education Singapore (MOE2017-T2-1-038, 1T1-02/04, 2017-T1-001-220 to Y.A.), (MOE2018-T2-1-085 to B.O.B.) and the Lee Kong Chian School of Medicine, Nanyang Technological University Start-up Grant (Y.A. and P.O.B.). This work is also partly supported by the LKCMedicine Healthcare Research Fund (Diabetes Research), established through the generous support of alumni of Nanyang Technological University. We thank Singhealth Transplant, the Singapore National Organ Transplant Unit and the Lee Foundation Grant for support in isolating human islets. D.G. is additionally supported by the Interdisciplinary Graduate Scholarship, Nanyang Technological University, Singapore. S.T.L is supported by the Nanyang President’s Graduate Scholarship.

## Materials and methods

### Animals

Homozygous diabetic (db/db) mice and nondiabetic control littermates of the B6.BKS(D)-Leprdb/J mouse strain were purchased from the Jackson Laboratory (Bar Harbor, ME) and maintained on a 12-hour light/dark cycle with free access to food and water. The CD169-DTR transgenic line was generated as described previously(51). Mice were strictly age-and sex-matched within experiments and were handled in accordance with institutional and national guidelines. With the exception of C57BL/6J mice which were purchased (InVivos Pte Ltd, Singapore), all other lines were kept as breeding colonies in local metabolic evaluation facilities under non-SPF conditions. All animal experiments were done with approval from institutional and university ethics committees (#140905, #A0373, #2013/SHS/816 and #2018/SHS/1399). To induce pre-diabetes, mice were either fed with high-fat diet (#D12492, Research Diets Inc, NJ, USA) or the calorie-matched control low-fat diet (#D12450J, Research Diets Inc, NJ, USA), accordingly. Diets were stored at −80°C and thawed to 4°C, for at least 24h before provided to animals *ad libitum*.

### Macrophage depletion by clodronate treatment

To analyze the role of macrophages on pancreatic islet function and insulin secretion during compensation, we have performed in vivo experiments to deplete macrophages from islets. Briefly, clodronate or PBS containing liposomes were administered intraperitoneally to 10 weeks old db/db or control mice (200 µL; 50mg/kg body weight; clodronateliposomes.com) and continued every 3 days, for 2weeks. Liposomes contained either PBS or clodronate, a non-toxic bisphosphonate. After injection, liposomes are ingested and digested by macrophages. Chlorinate is released and accumulates intracellular, where at high intracellular concentrations, it induces apoptosis(52). Animals were sacrificed at 12 weeks and a last dose of clodronate was given one day before the sacrifice. Animals were fasted overnight and intraperitoneal glucose tolerance test was performed. Blood was collected from tail vein for determining the plasma insulin levels. Pancreas excised and islets were isolated by collagenase digestion and used for functional analysis.

### Macrophage depletion in CD169-DTR mice

A total of 23 female C57BL/6 mice aged 8-9 weeks were used in this study, out of which 12 mice were CD169-DTR positive (DTR+) and 11 were DTR negative (DTR-) control mice.. All mice were fed with 60% HFD for 21 days. A total of 20 µg/ kg of DT was injected intraperitoneally every 3 days and subsequently the mice were culled to harvest pancreas and liver. The tissues harvested were fixed in 4% PFA and then cryopreserved for immunofluorescence studies.

### Macrophage depletion by anti-mouse CSF1R injection

C57BL/6 mice aged 10 weeks were were fed with 60% HFD for 21 days. 2mg of anti-mouse CSF1R (CD115, AFS98) (Bio X Cell, USA) antibody or PBS control was injected three times over 21 days (days 0, 10, and 20). The tissues were fixed in 4% paraformaldehyde for immunofluorescence studies. IPGTT, ITT and plasma insulin levels were measured on day0, 10 and 20.

### Glucose Tolerance Test, Insulin Tolerance Test, Insulin Secretion Test and insulin release

For glucose tolerance tests (GTTs), mice were fasted for 6–12 h. Next, mice were injected i.p with 2.0 g/kg.body weight of glucose and 1 U/kg body weight of insulin after 4h starvation for the insulin tolerance test (ITT). Before and up to 120 min after injection, blood glucose concentrations were measured in venous blood (Vena Saphena) using an automated glucometer. For the insulin secretion test (IST), blood samples were collected at 0, 15, and 30 min after glucose injection (2 g of glucose /kg body weight) for insulin determination by ELISA. The islets were individually dissected under a stereomicroscope. Batches of 10 islets of similar size were collected and incubated in RPMI 1640-10% fetal calf serum at 37°C in 5% CO_2_ for 2 h. These islets were washed and pre incubated in 0.5% (wt/vol) bovine serum albumin-Krebs-Ringer HEPES-buffered saline in 2.8 mM glucose at 37°C in 5% CO_2_ for 30 min and then transferred to 0.5% (wt/vol) bovine serum albumin-Krebs-Ringer HEPES-buffered saline in 2.8 mM glucose, stimulatory 11 mM glucose alone, or 25 mM KCl at 2.8 mM glucose. After incubation at 37°C in 5% CO_2_ for 30 min, the supernatants were measured for insulin release as described above.

### Immunolabelling

Pancreata were dissected and fixed overnight in 4% paraformaldehyde at 4°C, and placed in a 30% sucrose solution overnight at 4°C. Pancreata were embedded in O.C.T. (Tissue-Tek) and serial sectioned on a cryostat at 10mM. Indirect protein localization was obtained by incubations with primary antibodies overnight at 4°C. Sections were stained using guinea pig anti-insulin (1:200; Dako, CA, USA), rat anti-mouse F4/80, rat anti-mouse CD68 (1:200; Bio-Rad, CA, USA), rabbit anti-IBA-1 (1:200; Wako, VA, USA), goat anti-mouse CD31 (R&D Systems, MN, USA), rabbit anti-mouse Ki67 (abcam, MA, USA) and secondary anti-body conjugated to Alexafluor 488, 568 and 647 (1:400). Percentage of CD68^+^ and F4/80^+^ cells were determined by counting positive cells/total cells in the individual islet. Imaging and quantification was performed using Zeiss LSM 800 with airyscan confocal laser scanning microscope (Zeiss,Thornwood, NY 10594 United States) controlled with ZEN Blue image analyzer software and image J.

### Flow cytometry

16 weeks old lean and db/db lslets were sacrificed and the pancreatic islets were isolated. Briefly, islets were isolated from the pancreas of donor mice by collagenase digestion (0.8 mg/ml) (Sigma-Aldrich) followed by hand picking. Islets were washed with islet medium consisting of: RPMI-1640 medium supplemented with 1% (vol./vol.) l-glutamine, 1% (vol./vol.) penicillin– streptomycin, 11 mmol/l glucose and 10% (vol./vol.) FCS. Islets were then dissociated in to single cells by Accutase® (Thermo Fisher Scientific, MA, USA) digestion. Immunostaining and flow cytometry analyses were then performed according to standard procedures. The following antibodies were used: PE-Cy7-labelled CD45, APC-Cy7-labelled CD11b, PE-labelled F4/80, BV421-labelled MHC II and BV605-labelled Ly6C (Biolegend, San Diego, CA). Cells were pretreated with Fc block (clone 2.4G2). Single cell suspensions of enzyme dissociated pancreatic islets were stained for 30 min on ice and subsequently analyzed on a Fortessa X-20 flow cytometer (BD Biosciences, CA, USA).

### Cytokine/chemokine analysis *ex vivo*

Mouse Islets were isolated as described above. Isolated islets (≈100) were cultured overnight in CMRL-1066 medium containing 10% FCS, 2mmol/L L-glutamine, 100 units/mL penicillin, and 0.1mg/mL streptomycin (Complete CMRL). Cytokine/chemokines in mouse islet culture supernatant were detected utilizing the Milliplex® MAP Mouse Cytokine/Chemokine Magnetic Bead Panel (EMD Millipore Corporation, Billerica, MA) according to manufacturer instructions. Briefly, supernatant was plated on 96-well plates. Following overnight incubation with magnetic beads, samples were washed and incubated at room temperature with detection antibodies. Samples were detected following addition of streptavidin-phycoerythrin on a Bio-Plex 200 and analyzed Bio-Plex manager™ Software (Bio-Rad, CA). Human islet preparations used in our study was in accordance ethical approval (NTU Singapore, #IRB-2018-05-052) and to reporting regulations as described previously (53). HP-006, HP-007 and HP-18330-01 are unique identifiers of human islets isolated from females aged 46, 58 and 53, respectively and with no history of diabetes (HbA1c <6.0%). Islets were obtained either in-house (HP-006, HP-007) or from OneLegacy (HP-18330-01). Isolated human islets were cultured overnight in CMRL-1066 media as described before. Islets were then treated ex vivo with liposomes containing PBS or clodronate (200µg/mL) for 16 hrs. Islets were washed replenished with fresh media, incubated further overnight. Supernatants were collected for cytokine/growth factor analysis. Islets were used for immunofluorescence (to confirm macrophage depletion) and qPCR. TNF-A, IL-10 and IL-6 were measured using Milliplex® MAP human Cytokine/Chemokine Magnetic Bead Panel. hVEGF was analyzed using DuoSet ELISA development system (R&D Systems, Inc. MN, USA) according to manufacturer’s instructions.

### Islet isolation and transplantation into the anterior chamber of the eye (ACE)

12 weeks old C57BL/6 mice (N=10) were fed with HFD for 21 days. After 21 days, 4 donor animals were sacrificed and the pancreatic islets were isolated as described above. Islets were then left to rest overnight at 37°C and 5% CO2, in CMRL1066 media. Islets were treated with liposome containing clodronate or its control for 24h hr and washed. Recipient HFD-fed mice were anesthetized by inhalation of ∼2% isoflurane in 30 oxygen via a face mask. A 23-gauge needle was used to make a small incision in the cornea, close to the corneal limbus, and 10 islets in phosphate-buffered saline (PBS) were slowly injected into the ACE using a 25G blunt cannula (vendor), making sure the islets were distributed throughout the surface area of the iris to favor their engraftment. Islets were allowed to engraft for 4 weeks.

### In vivo imaging of islets transplanted to the anterior chamber of the eye

At the indicated time points after transplantation (see Results and figure legends), we anesthetized mice with a 40% oxygen and a ∼2.5% isoflurane mixture. Animal positioning and preparation for imaging was performed as described before (54). For imaging, we used an upright DM6000 Leica microscope equipped with a TCS-SP8-AOBS confocal scanner (Leica Microsystems). Viscotears (Novartis) was used as immersion liquid. To visualize the islet (and iris) vessels, the animal was injected with 150 KDa FITC-labelled dextran (50 mg/kg, Sigma, St. Louis, MO). Leica Confocal Software (version 2.61), and ImageJ were used to process images.

### Endothelial tube formation (Angiogenesis) assay

Human pancreatic microvascular endothelial cells (HPaMEC, ScienCell, USA) were maintained in endothelial cell medium (ECM) supplemented with 5% fetal bovine serum, 1% endothelial cell growth supplement (ECGS) and 1% penicillin/streptomycin according to supplier’s instructions. 50μL of Growth Factor Reduced Matrigel Basement Membrane Matrix (Corning, USA) was added in each well of a 96-well plate and incubated at 37°C for 30 minutes. 20,000 HPaMECs were resuspended in 40μL of ECM and 40μL of the respective conditioned media in a 1:1 ratio and seeded onto the polymerized Matrigel. After a 22-hour incubation, the vasculature was visualised using Eclipse Ti-E Inverted Research Microscope (Nikon, USA) and the complexity of the vascular network was analyzed by Image J (National Institutes of Health, USA). Duplicates were performed for each treatment.

### Statistics

Data collected is assumed to be normally distributed and are therefore expressed as mean ± SEM, with a minimum of 3 independent experiments each. Statistical significance was calculated by two-tailed Student’s t-test, or by ANOVA, as detailed in each figure legend. *P*≤ 0.05 is considered as statistically significant.

## Supplemental figure legends

**Supp. Figure 1.**
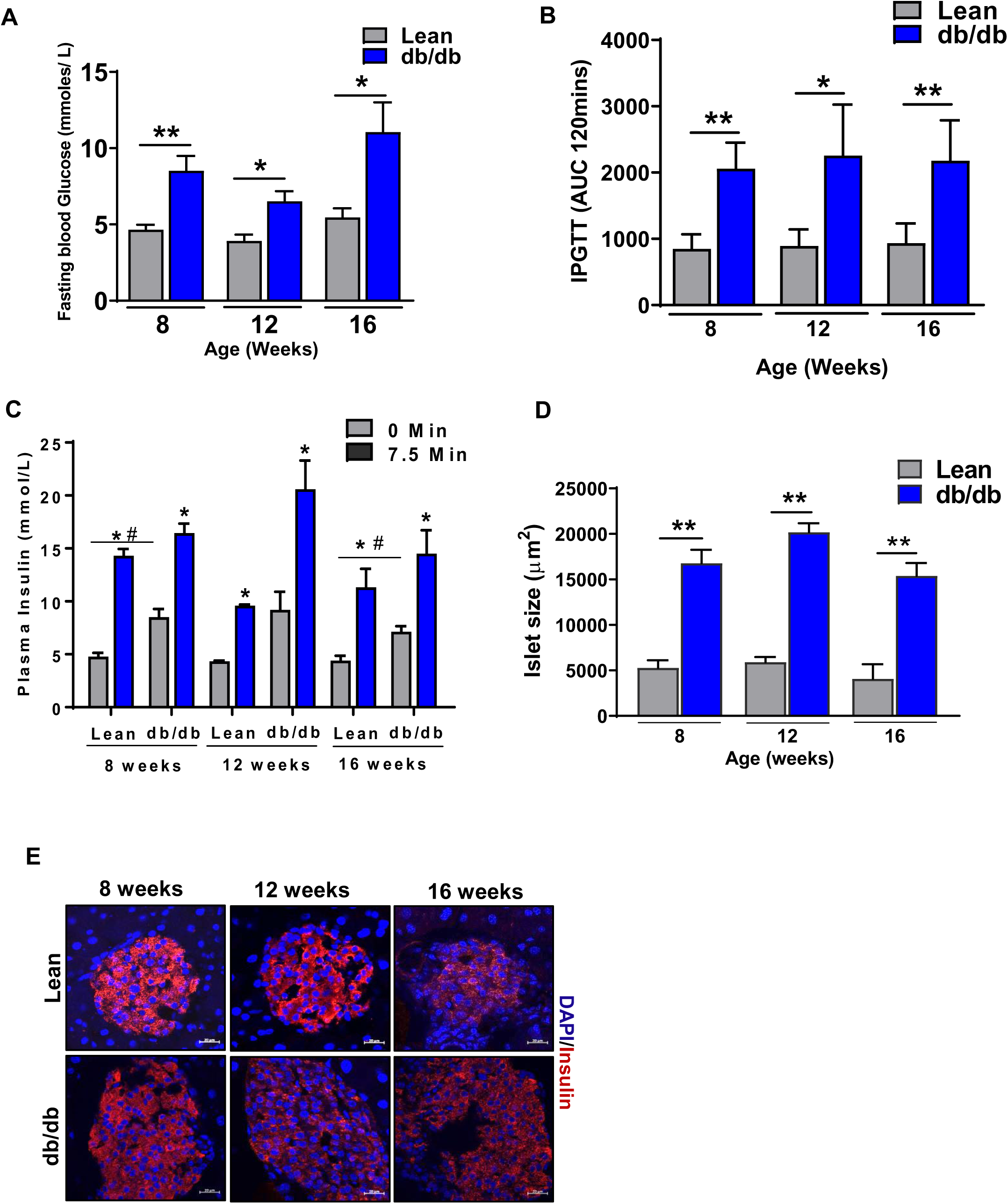
db/db mouse is a model of compensatory diabetes before the age of 16 weeks. **A.** Fasting blood glucose levels in db/db and age matched lean controls of different ages. **B.** Glucose tolerance tests (IPGTT) after intraperitoneal loading with 2 g D-glucose/kg were performed on db/db and age matched lean controls of different ages after overnight fasting. Glucose tolerance is expressed as area under the curve (AUC). **C.** Level of plasma insulin before and after intraperitoneal loading with 2 g D-glucose/kg were performed on db/db and age matched lean controls of different ages after overnight fasting. **D.** Increased Islet size in db/db mice at different ages. **E.** Representative fluorescent images; Insulin (red), DAPI (blue). All data in this figure are the means ± SEM (scale bar, 20μm). p-values were calculated using Student’s t-test *p<0.05, **p<0.005, ^#^p< 0.05 Vs lean.

**Supp. Figure 2.**
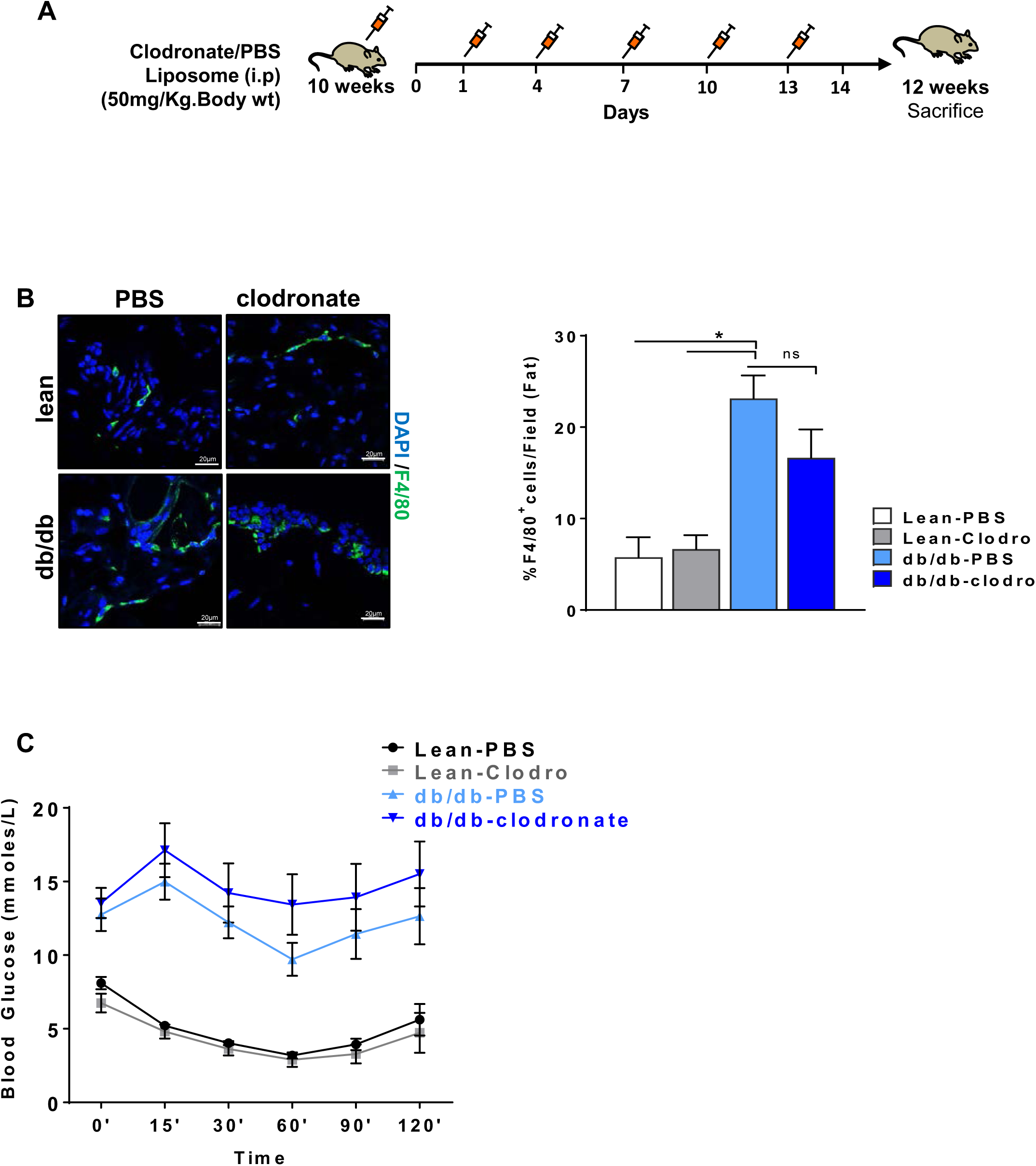
No significant change in peripheral insulin sensitivity after 2-week clodronate treatment. **A.** Schematic representation of the experimental design. **B.** Representative Immunofluorescence image showing depletion of F4/80^+^ macrophages after clodronate treatment in adipose tissue (left panel) and quantification of F4/80^+^ macrophages/field (Right panel). F4/80 (green), DAPI (blue)**. C.** Insulin tolerance test done by 4hr fasting followed by i.p. injection of insulin (1.0 U kg^−1^) and measurement of blood glucose concentrations at indicated time intervals. All data in this figure are the means ± SEM (scale bar, 20 μm). p-values were calculated using One Way ANOVA with Tukey’s multiple comparisons test (**b**) and unpaired Student’s t-test (**c**) *P<0.05 Vs lean.

**Supp. Figure 3.**
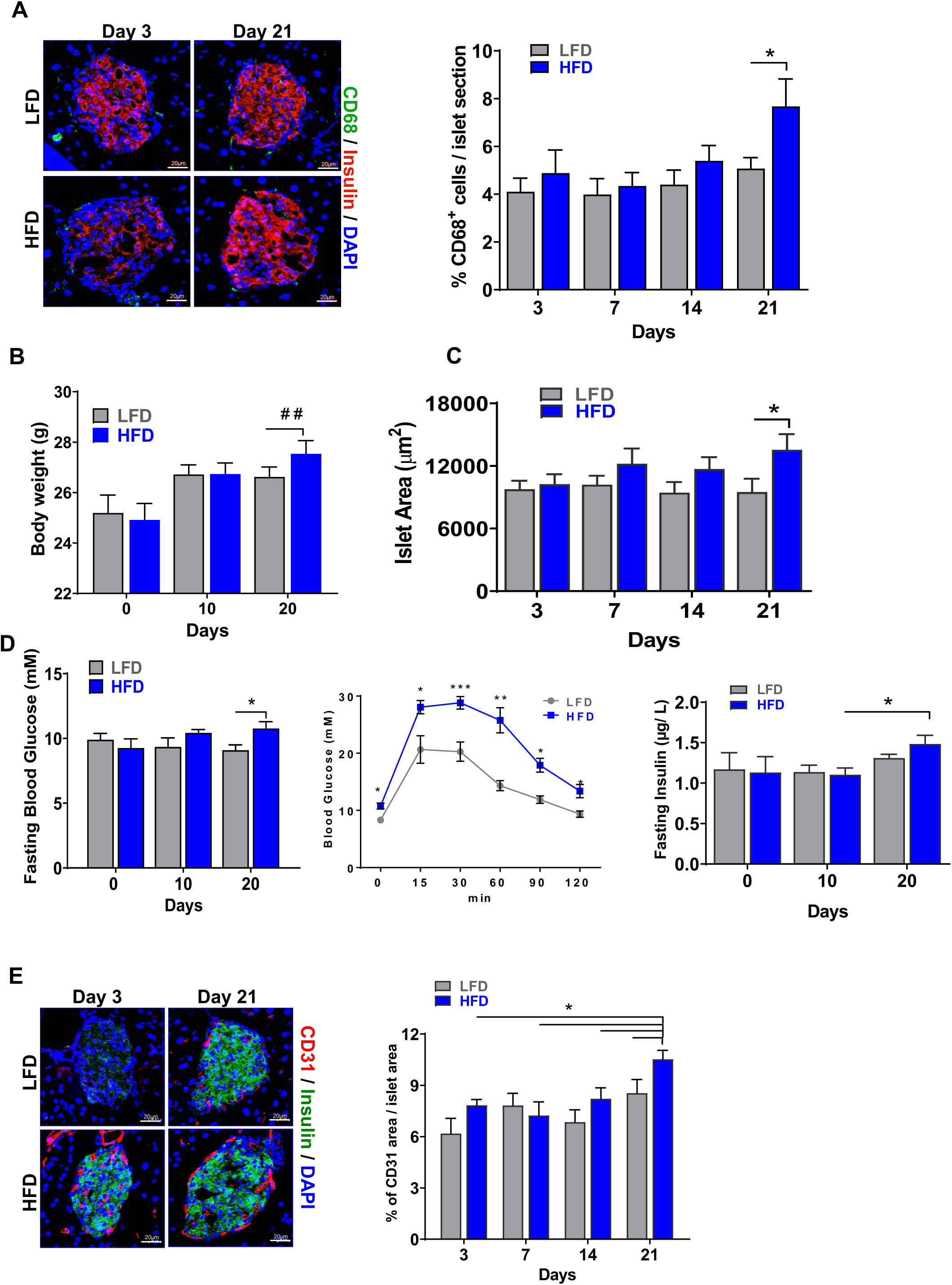
Metabolic phenotype of acute HFD fed mice. **A.** Representative image showing CD68^+^ macrophages in HFD fed mice compared to LFD fed mice (left panel) and quantification of CD68^+^ cells/islet (right panel). Insulin (red), CD68 (green), DAPI (blue). **B.** Body weight. **C.** Islet area. **D.** Fasting blood glucose (left panel), IPGTT (middle panel), Fasting Insulin (right panel). **E.** Representative image (left panel) showing effect of acute HFD feeding on islet vasculature. Insulin (green), endothelial marker CD31 (red) and DAPI (blue). Quantitation of blood vessel area (right panel). All data in this figure are the means ± SEM (scale bar, 20 μm). P-values were calculated using unpaired Student’s t-test ***p<0.001, **p<0.005 *p<0.05 Vs LFD.

**Supp. Figure 4.**
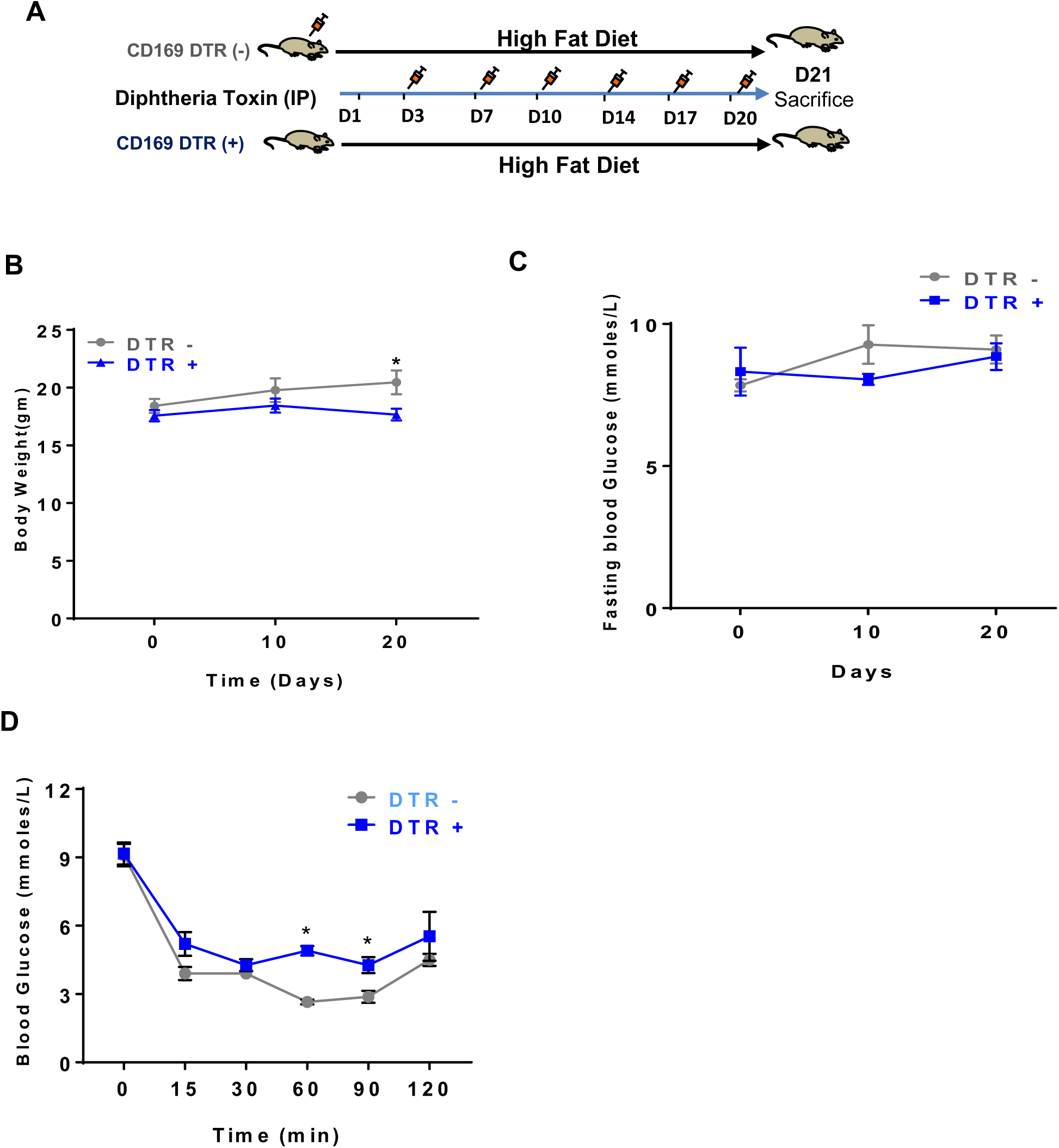
Changes in metabolic phenotype upon macrophage ablation in CD169-DTR mice. **A.** Schematic representation of the experimental design for macrophage depletion in CD169-DTR^+^ mouse model. **B.** Body weight. **C.** Fasting Blood glucose. **D.** Insulin tolerance test (ITT) after 4hr fasting. All data in this figure are the means ± SEM. P-values were calculated using unpaired Student’s t-test *p<0.05.

**Supp. Figure 5.**
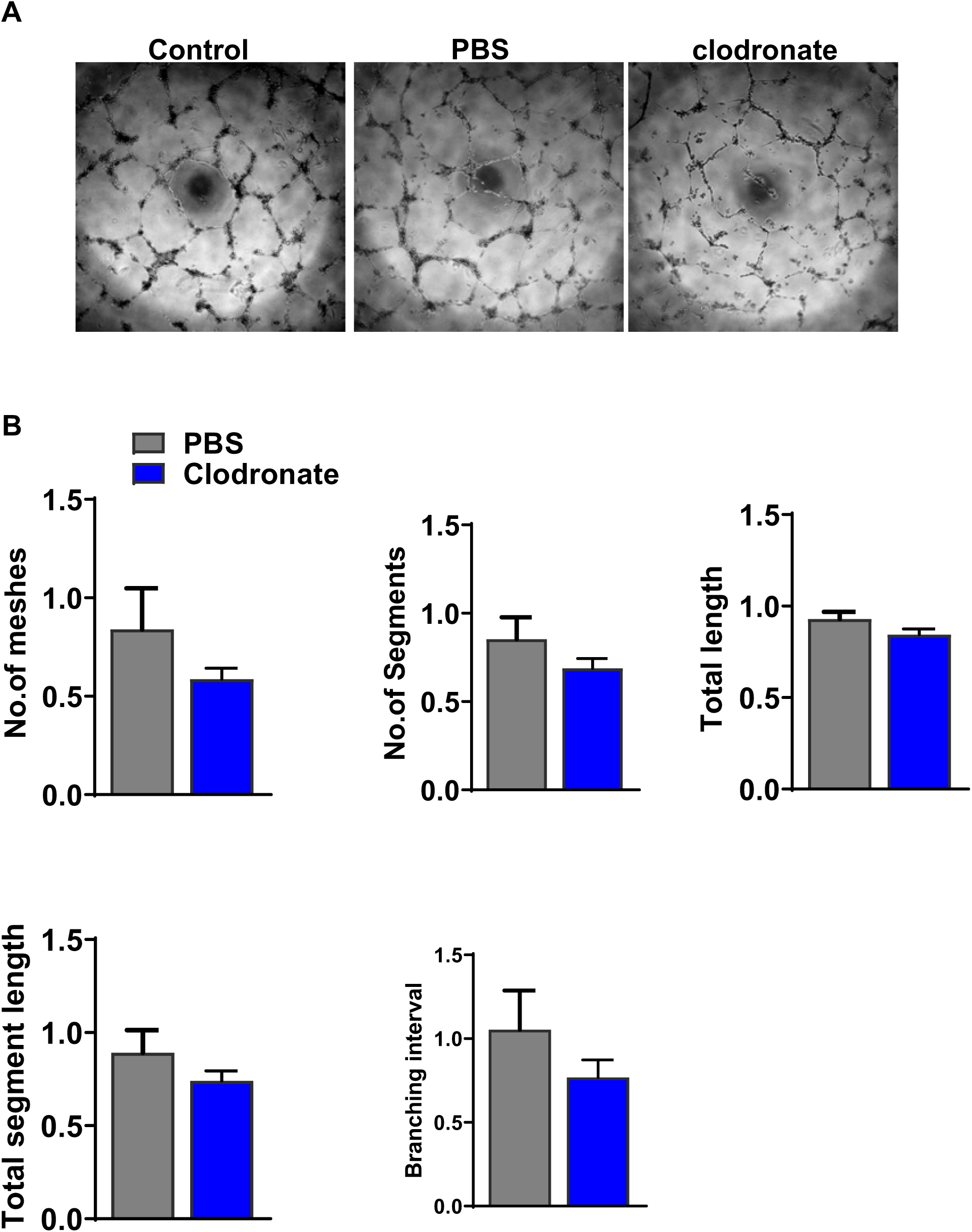
Supernatants from macrophage-depleted human islets disrupts human pancreatic endothelial tube formation *in vitro* **A**. Representative images (left) of HPaMEC tube formation in growth factor reduced Matrigel upon treatment with respective conditioned media. **B.** Quantification of the number of meshes, number of segments, total tube length, total segment length and branching interval. Data are presented as the mean ± SEM; n= 3

## References

1. Kahn SE HR, Utzschneider KM. Mechanisms linking obesity to insulin resistance and type 2 diabetes. Nature. 2006;444:840–6

2. Lackey DE, Olefsky, J.M.,. Regulation of metabolism by the innate immune system. Nature Reviews Endocrinology. 2016;12(1):15–28.

3. Ying W, Lee YS, Dong Y, Seidman JS, Yang M, Isaac R, et al. Expansion of Islet-Resident Macrophages Leads to Inflammation Affecting beta Cell Proliferation and Function in Obesity. Cell Metab. 2018.

4. Kammoun HL, Kraakman M.J., Febbraio, M.A.,. Adipose tissue inflammation in glucose metabolism. Reviews in Endocrine and metabolic disorders 2014;15(1):31–44.

5. Okabe Y, Medzhitov, R.,. Tissue-specific signals control reversible program of localization and functional polarization of macrophages. Cell. 2014;157(4):832–44

6. Kraakman MJ MA, Jandeleit-Dahm K, Kammoun HL. Macrophage polarization in obesity and type 2 diabetes: weighing down our understanding of macrophage function?. Front Immunol. 2014;5(470):1–6.

7. Osborn O OJ. The cellular and signaling networks linking the immune system and metabolism in disease. Nat Med. 2012;18(3):363–74

8. Xiao X GI, Guo P, Wiersch J, Fischbach S, Peirish L, Song Z, El-Gohary Y, Prasadan K, Shiota C, Gittes GK. M2macrophages promote beta-cell proliferation by up-regulation of SMAD7. Proc Natl Acad Sci U SA. 2014;111(13):E1211–E20.

9. Dinarello CA, Donath, M.Y., Mandrup-Poulsen, T.,. Role of IL-1β in type 2 diabetes. Current Openion in Endocrinology Diabetes and Obesity. 2010;17(4):314–21.

10. Donath MY, Böni-Schnetzler, M., Ellingsgaard, H., Ehses, J.A.,. Islet inflammation impairs the pancreatic β-cell in type 2 diabetes. Physiology. 2009;24(6):325–31.

11. Odegaard JI, Chawla, A.,. Connecting type 1 and type 2 diabetes through innate immunity. Cold Spring Harbor Perspective in Medicine. 2012;2(3):1–18.

12. Richardson SJ WA, Bone AJ, Foulis AK, Morgan NG. Islet-associated macrophages in type 2 diabetes. Diabetologia. 2009;52:1686–8

13. Kamata K MH, Inaba W, et al.; 21:191–201 (2014). Islet amyloid with macrophage migration correlates with augmented β-cell deficits in type 2 diabetic patients. Amyloid. 2014;21:191–201

14. Eguchi K MI, Oishi-Tanaka Y, Ohsugi M, Kono N, Ogata F, Yagi N, Ohto U, Kimoto M, Miyake K, Tobe K, Arai H, Kadowaki T, Nagai R. Saturated fatty acid and TLR signaling link β cell dysfunction and islet inflammation. Cell Metab. 2012;15:518–33

15. Jourdan T GG, Cinar R, Bertola A, Szanda G, Liu J, Tam J, Han T, Mukhopadhyay B, Skarulis MC, Ju C, Aouadi M, Czech MP, Kunos G. Activation of the Nlrp3 inflammasome in infiltrating macrophages by endocannabinoids mediates β cell loss in type 2 diabetes. Nat Med. 2013;19:1132–40

16. Masters SL DA, Subramanian SL, Hull RL, Tannahill GM, Sharp FA, Becker C, Franchi L, Yoshihara E, Chen Z, Mullooly N, Mielke LA, Harris J, Coll RC, Mills KH, Mok KH, Newsholme P, Nuñez G, Yodoi J, Kahn SE, Lavelle EC, O’Neill LA Activation of the NLRP3 inflammasome by islet amyloid polypeptide provides a mechanism for enhanced IL-1β in type 2 diabetes. Nat Immunol. 2010;11:897–904

17. Westwell-Roper C DD, Soukhatcheva G, Potter KJ, van Rooijen N, Ehses JA, Verchere CB. IL-1 blockade attenuates islet amyloid polypeptide-induced proinflammatory cytokine release and pancreatic islet graft dysfunction. J Immunol. 2011;187:2755–65

18. Nackiewicz D1 DM, He W, Kim R, Salmi A, Rütti S, Westwell-Roper C, Cunningham A, Speck M, Schuster-Klein C, Guardiola B, Maedler K, Ehses JA TLR2/6 and TLR4-activated macrophages contribute to islet inflammation and impair β cell insulin gene expression via IL-1 and IL-6. Diabetologia. 2014;57:1645–54.

19. Banaei-Bouchareb L1 G-EV, Samara-Boustani D, Castellotti MC, Czernichow P, Pollard JW, Polak M. Insulin cell mass is altered in Csf1op/Csf1op macrophage deficient mice. J Leukoc Biol. 2004;76:359–67

20. Criscimanna A CG, Gittes GK, Piganelli JD, Esni F. Activated macrophages create lineage-specific microenvironments for pancreatic acinar- and β-cell regeneration in mice. Gastroenterology. 2014;147:1106–18

21. Brissova M, Aamodt, K., Brahmachary, P., Prasad, N., Hong, J.Y., Dai, C., Mellati, M., Shostak, A., Poffenberger, G., Aramandla, R., Levy, S.E., Powers, A.C.,. Islet microenvironment, modulated by vascular endothelial growth factor-A signaling, promotes β cell regeneration. Cell Metabolism. 2014;19(3):498–511.

22. Morris DL. Minireview: Emerging Concepts in Islet Macrophage Biology in Type 2 Diabetes. Mol Endocrinol. 2015;29(7):946–62.

23. Calderon B, Carrero, J.A., Ferris, S.T., Sojka, D.K., Moore, L., Epelman, S., Murphy, K.M., Yokoyama, W.M., Randolph, G.J., Unanue, E.R.,. The pancreas anatomy conditions the origin and properties of resident macrophages. Journal of Experimental Medicine 2015; 212(10):1497–512

24. Do OH, Gunton JE, Gaisano HY, and Thorn P. Changes in beta cell function occur in prediabetes and early disease in the Lepr (db) mouse model of diabetes. Diabetologia. 2016;59(6):1222–30.

25. Chen C CCM, Stertmann J, Bozsak R, Speier S. Human beta cell mass and function in diabetes: Recent advances in knowledge and technologies to understand disease pathogenesis. Molecular Metabolism 6 2017;6(9):943– 57.

26. Nolan C.J PM. Islet beta cell failure in type 2 diabetes. Journal of Clinical Investigation. 2006;116(7):1802–12.

27. Agudo J AE, Jimenez V, Casellas A, Mallol C, Salavert A, Tafuro S, Obach M, Ruzo A, Moya M, Pujol A, Bosch F. Vascular endothelial growth factor-mediated islet hypervascularization and inflammation contribute to progressive reduction of β-cell mass. Diabetes. 2012;61(11):2851–61.

28. Brissova M SA, Shiota M, Wiebe P.O, Poffenberger G, Kantz J, Chen Z, Carr C, Jerome WG, Chen J, Baldwin HS, Nicholson W, Bader DM, Jetton T, Gannon M, Powers AC. Pancreatic Islet Production of Vascular Endothelial Growth Factor-A Is Essential for Islet Vascularization, Revascularization, and Function. Diabetes. 2006;55(11):2974–85.

29. Cai Q BM, Reinert RB, Pan FC, Brahmachary P, Jeansson M, Shostak A, Radhika A, Poffenberger G, Quaggin SE, Jerome WG, Dumont DJ, Powers AC. Enhanced expression of VEGF-A in β cells increases endothelial cell number but impairs islet morphogenesis and β cell proliferation. Dev Biol. 2012;367(1):40–54.

30. Ferris ST, Zakharov PN, Wan X, Calderon B, Artyomov MN, Unanue ER, et al. The islet-resident macrophage is in an inflammatory state and senses microbial products in blood. J Exp Med. 2017;214(8):2369–85.

31. Ehses JA, Perren, A., Eppler, E., Ribaux, P., Pospisilik, J.A., Maor-Cahn, R., Gueripel, X., Ellingsgaard, H., Schneider, M.K., Biollaz, G., Fontana, A., Reinecke, M., Homo-Delarche, F., Donath, M.Y.,. Increased number of islet-associated macrophages in Type 2 diabetes. Diabetes. 2007;56(9):2356–70

32. Campbell IL, Cutri A, Wilson A, and Harrison LC. Evidence for IL-6 production by and effects on the pancreatic beta-cell. J Immunol. 1989;143(4):1188–91.

33. Woodland DC, Liu W, Leong J, Sears ML, Luo P, and Chen X. Short-term high-fat feeding induces islet macrophage infiltration and beta-cell replication independently of insulin resistance in mice. Am J Physiol Endocrinol Metab. 2016;311(4):E763–E71.

34. Gupta P LS, Sheng J, Tetlak P, Balachander A, Claser C, Renia L, Karjalainen K, Ruedl C. Tissue-Resident CD169(+) Macrophages Form a Crucial Front Line against Plasmodium Infection. Cell Reports. 2016;16(6):1749–61.

35. Bu L, Gao M, Qu S, and Liu D. Intraperitoneal injection of clodronate liposomes eliminates visceral adipose macrophages and blocks high-fat diet-induced weight gain and development of insulin resistance. AAPS J. 2013;15(4):1001–11.

36. Carrero JA MD, Ferris ST, Wan X, Hu H, Zinselmeyer BH, Vomund AN, Unanue ER. Resident macrophages of pancreatic islets have a seminal role in the initiation of autoimmune diabetes of NOD mice. Proc Natl Acad Sci USA. 2017;114(48):E10418–E27.

37. Almaça J MJ, Arrojo E Drigo R, Abdulreda MH, Jeon WB, Berggren PO, Caicedo A, Nam HG. Young capillary vessels rejuvenate aged pancreatic islets. Proc Natl Acad Sci USA. 2014;111(49):17612–7.

38. Speier S ND, Köhler M, Caicedo A, Leibiger IB, Berggren PO. Noninvasive high-resolution in vivo imaging of cell biology in the anterior chamber of the mouse eye. Nat Protocols. 2008;3(8):1278–86.

39. Diez JA AEDR, Zheng X, Stelmashenko OV, Chua M, Rodriguez-Diaz R, Fukuda M, Köhler M, Leibiger I, Tun SBB, Ali Y, Augustine GJ, Barathi VA, Berggren PO. Pancreatic Islet Blood Flow Dynamics in Primates. Cell reports. 2017;20(6):1490–501.

40. Susan J. Burke HMB, David H. Burk, Robert C. Noland, Adrianna E. Eder, Matthew S. Boulos, Michael D. Karlstad, J. Jason Collier. db/db Mice Exhibit Features of Human Type 2 Diabetes That Are Not Present in Weight-Matched C57BL/6J Mice Fed a Western Diet. J Diabetes Res. 2017;2017:1–17.

41. Dalboge LS, Almholt DL, Neerup TS, Vassiliadis E, Vrang N, Pedersen L, et al. Characterisation of age-dependent beta cell dynamics in the male db/db mice. PLoS One. 2013;8(12):e82813.

42. Linnemann AK, Blumer J, Marasco MR, Battiola TJ, Umhoefer HM, Han JY, et al. Interleukin 6 protects pancreatic beta cells from apoptosis by stimulation of autophagy. FASEB J. 2017;31(9):4140–52.

43. Choi SE, Choi KM, Yoon IH, Shin JY, Kim JS, Park WY, et al. IL-6 protects pancreatic islet beta cells from pro-inflammatory cytokines-induced cell death and functional impairment in vitro and in vivo. Transpl Immunol. 2004;13(1):43–53.

44. Suzuki T, Imai J, Yamada T, Ishigaki Y, Kaneko K, Uno K, et al. Interleukin-6 enhances glucose-stimulated insulin secretion from pancreatic beta-cells: potential involvement of the PLC-IP3-dependent pathway. Diabetes. 2011;60(2):537–47.

45. Johnson AR, Milner, J.J, Makowski, L.). The inflammation highway: metabolism accelerates inflammatory traffic in obesity. Immunol Rev 2012;249: 218–38

46. Donath MY B-SM, Ellingsgaard H, Ehses JA. Islet inflammation impairs the pancreatic β-cell in type 2 diabetes. Physiology. 2009;24:325–33.

47. Boni-Schnetzler M TJ, Parnaud G, Marselli L, Ehses JA, Kerr-Conte J, Pattou F, Halban PA, Weir GC, Donath MY. Increased interleukin (IL)-1beta messenger ribonucleic acid expression in beta-cells of individuals with Type 2 diabetes and regulation of IL-1beta in human islets by glucose and autostimulation J Clin Endocrinol Metab. 2008; 93: 4065–74

48. Ehses JA PA, Eppler E, Ribaux P, Pospisilik JA, Maor-Cahn R, Gueripel X, Ellingsgaard H, Schneider MK, Biollaz G, Fontana A, Reinecke M, Homo-Delarche F, Donath MY Increased number of islet-associated macrophages in Type 2 diabetes. Diabetes 2007;56:2356–70

49. Riley KG PR, Maulis MF, Dunn JC, Bolus WR, Kendall PL, Hasty AH, Gannon M. Macrophages are essential for CTGF-mediated adult β-cell proliferation after injury. Molecular Metabolism 2015;4(8):584–91.

50. Jonathan R. Weitz MM, Joana Almaça, Julia Stertmann, Kristie Aamodt, Marcela Brissova, Stephan Speier, Rayner Rodriguez-Diaz, Alejandro Caicedo. Mouse pancreatic islet macrophages use locally released ATP to monitor beta cell activity. Diabetologia. 2018;61(1):182–92.

51. Purnama C, Ng SL, Tetlak P, Setiagani YA, Kandasamy M, Baalasubramanian S, et al. Transient ablation of alveolar macrophages leads to massive pathology of influenza infection without affecting cellular adaptive immunity. Eur J Immunol. 2014;44(7):2003–12.

52. Van Rooijen N, Hendrikx, E.,. Liposomes for Specific Depletion of Macrophages from Organs and Tissues. Methods in Molecular Biology. 2010;605:189–203.

53. Powers NJHC. Use of human islets to understand islet biology and diabetes: progress, challenges and suggestions. Diabetologia. 2019;62(2):212–2222.

54. Stephan Speier DN, Over Cabrera, Jia Yu, R Damaris Molano, Antonello Pileggi, Tilo Moede, Martin Köhler, Johannes Wilbertz3, Barbara Leibiger, Camillo Ricordi, Ingo B Leibiger, Alejandro Caicedo & Per-Olof Berggren. Noninvasive in vivo imaging of pancreatic islet cell biology. Nature Medicine. 2008;14(5):574–8.

